# Molecular structure of a prevalent amyloid-β fibril polymorph from Alzheimer’s disease brain tissue

**DOI:** 10.1101/2020.03.06.981381

**Authors:** Ujjayini Ghosh, Kent R. Thurber, Wai-Ming Yau, Robert Tycko

## Abstract

Amyloid-β (Aβ) fibrils exhibit self-propagating, molecular-level polymorphisms that may underlie variations in clinical and pathological characteristics of Alzheimer’s disease. We report the molecular structure of a specific brain-derived polymorph that has been identified as the most prevalent polymorph of 40-residue Aβ fibrils in cortical tissue of Alzheimer’s disease patients. This structure, developed from cryo-electron microscopy and supported by solid state NMR data, differs qualitatively from all previously described Aβ fibril structures, both in its molecular conformation and its organization of cross-β subunits. Knowledge of this brain-derived fibril structure may contribute to the development of structure-specific amyloid imaging agents and aggregation inhibitors with greater diagnostic and therapeutic utility.

## Main Text

Fibrillar assemblies formed by amyloid-β (Aβ) peptides are the primary proteinaceous component of the amyloid plaques, diffuse amyloid, and vascular amyloid in brain tissue of Alzheimer’s disease (AD) patients. The most abundant forms of Aβ are 40 and 42 amino acids in length (Aβ40 and Aβ42). Mature fibrils formed by full-length Aβ peptides always contain in-register, parallel β-sheets with a cross-β orientation relative to the fibril growth direction (*1–3*). However, studies of Aβ fibrils grown *in vitro* have shown that other aspects of their molecular structures are variable (*4–10*). Moreover, when new Aβ fibrils are grown from fragments of existing fibrils (*i.e.*, from fibril seeds), molecular structural differences are preserved. Thus, Aβ fibrils exhibit self-propagating, molecular-level polymorphism (*4*). Full molecular structural models for various Aβ40 and Aβ42 fibril polymorphs prepared *in vitro* have been developed from solid state nuclear magnetic resonance (ssNMR) data (*6, 7, 11–16*) and from cryogenic electron microscopy (cryoEM) (*17*).

Evidence that structural variations in Aβ fibrils may correlate with variations in clinical and pathological characteristics of AD (*18–23*) provides a strong motivation for detailed structural characterization of brain-derived fibril polymorphs. Spectroscopic signatures (*i.e.*, NMR chemical shifts) of the most common Aβ40 and Aβ42 fibril polymorphs in AD brain were identified in recent ssNMR measurements by Qiang *et al*. (*24*), in which isotopically labeled Aβ fibrils were prepared by seeded growth from cerebral cortical tissue. Based on these and other studies (*25, 26*), it appears that Aβ fibril polymorphs in human brain tissue are different from those that have been characterized *in vitro*. Here we report the molecular structure of the Aβ40 fibril polymorph that is the most common polymorph in AD brain (*24*).

Aβ40 fibrils for structural measurements by cryoEM and ssNMR were prepared by seeding solutions of synthetic or recombinant Aβ40 with fibrils that had been produced by seeded growth from cortical tissue extracts from two AD patients (*27*). These samples are therefore “second-generation” fibrils. Negative-stain transmission electron microscope (TEM) and cryoEM images (Fig. 1A,B) show that the fibrils exhibit the same modulation of their apparent widths as the first-generation, brain-seeded fibrils from which they derive (*24, 25*), with crossover distances (i.e., distances between width minima) of about 120 nm (Fig. S1A,B). Second-generation and first-generation fibrils also have the same ssNMR spectra (*28*). Importantly, the fact that second-generation fibril samples contain less extraneous material from the original brain extracts allowed us to obtain high-quality cryoEM images, suitable for molecular structure determination.

**Figure 1:**
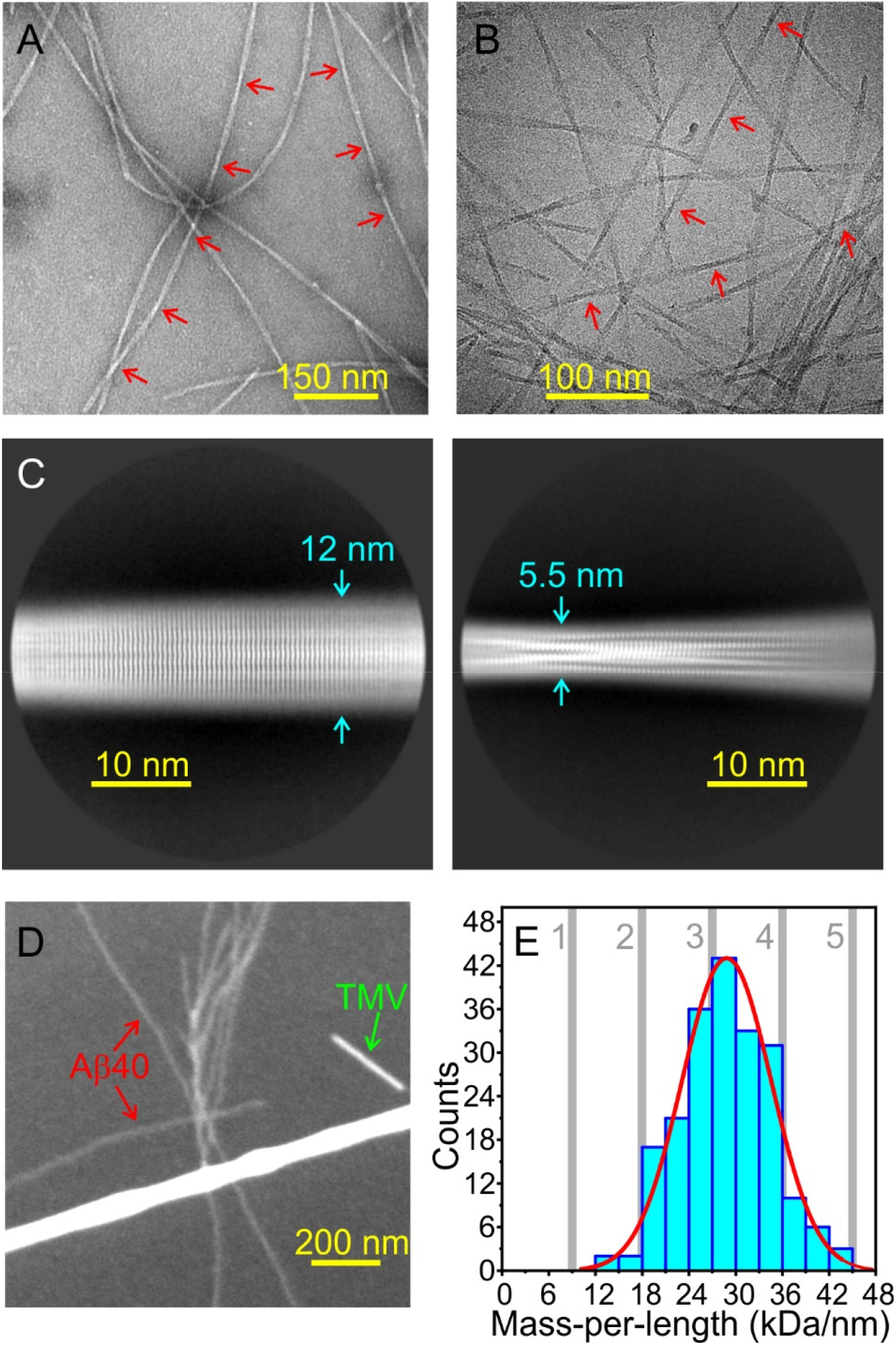
Morphology of brain-derived Aβ40 fibrils. (A) Negative-stain TEM image of second-generation fibrils used for cryoEM. Red arrows indicate approximate crossover points. (B) Example of cryoEM image, low-pass filtered to make fibrils more readily visible. (C) Examples of 2D classes, generated from 241,105 particles in 1337 cryoEM images. (D) Example of dark-field TEM image of unstained third-generation fibrils. TMV rods serve as intensity standards for quantification of fibril MPL. (E) MPL histogram from dark-field TEM images. Vertical gray bars indicate theoretical MPL values for 1-5 cross-β units. A Gaussian fit centered at 28.8 kDa/nm with 13.5 kDa/nm FWHM is shown in red.

Fig. 1C shows examples of two-dimensional (2D) class averages derived from cryoEM images. Pronounced striations perpendicular to the fibril growth direction, with a spacing of about 4.8 Å, indicate the expected cross-β structure. All 2D classes have approximate reflection symmetry with respect to their midline, indicating a structure with two-fold rotational or screw symmetry about the growth direction. However, measurements of the mass-per-length (MPL) of these fibrils, by quantification of intensities in dark-field TEM images (*5, 29*), yield MPL = 28.8 ± 0.4 kDa/nm (Figs. 1D,E and S1C,D). Given the 4.33 kDa molecular mass of Aβ40 and the 4.8 Å cross-β spacing, an Aβ40 fibril structure consisting of two cross-β subunits with the usual in-register, parallel β-sheet organization would have MPL ≈ 18 kDa/nm, while a structure consisting of three cross-β subunits would have MPL ≈ 27 kDa/nm (*4, 5, 7, 25, 29*). Thus, we encounter the conundrum of a structure that contains three cross-β subunits, somehow arranged with approximate two-fold symmetry about the growth direction.

To derive a three-dimensional (3D) density map from the cryoEM images, we used RELION software for helical reconstruction (*30, 31*), modified to include correlations of particle orientations about the fibril growth direction for particles from the same fibril segment (see Supporting Text). Calculations were performed without additional symmetry, with two-fold rotational symmetry in the repeat unit, and with 2_1_ screw symmetry. The density map with the highest final resolution (2.77 Å) was obtained with near 2_1_ symmetry, generated by a rise of 2.45 Å and twist of −180.34° between repeats (Fig. 2A). A density map without additional symmetry and Fourier-shell correlation plots are shown in Fig. S2.

**Figure 2:**
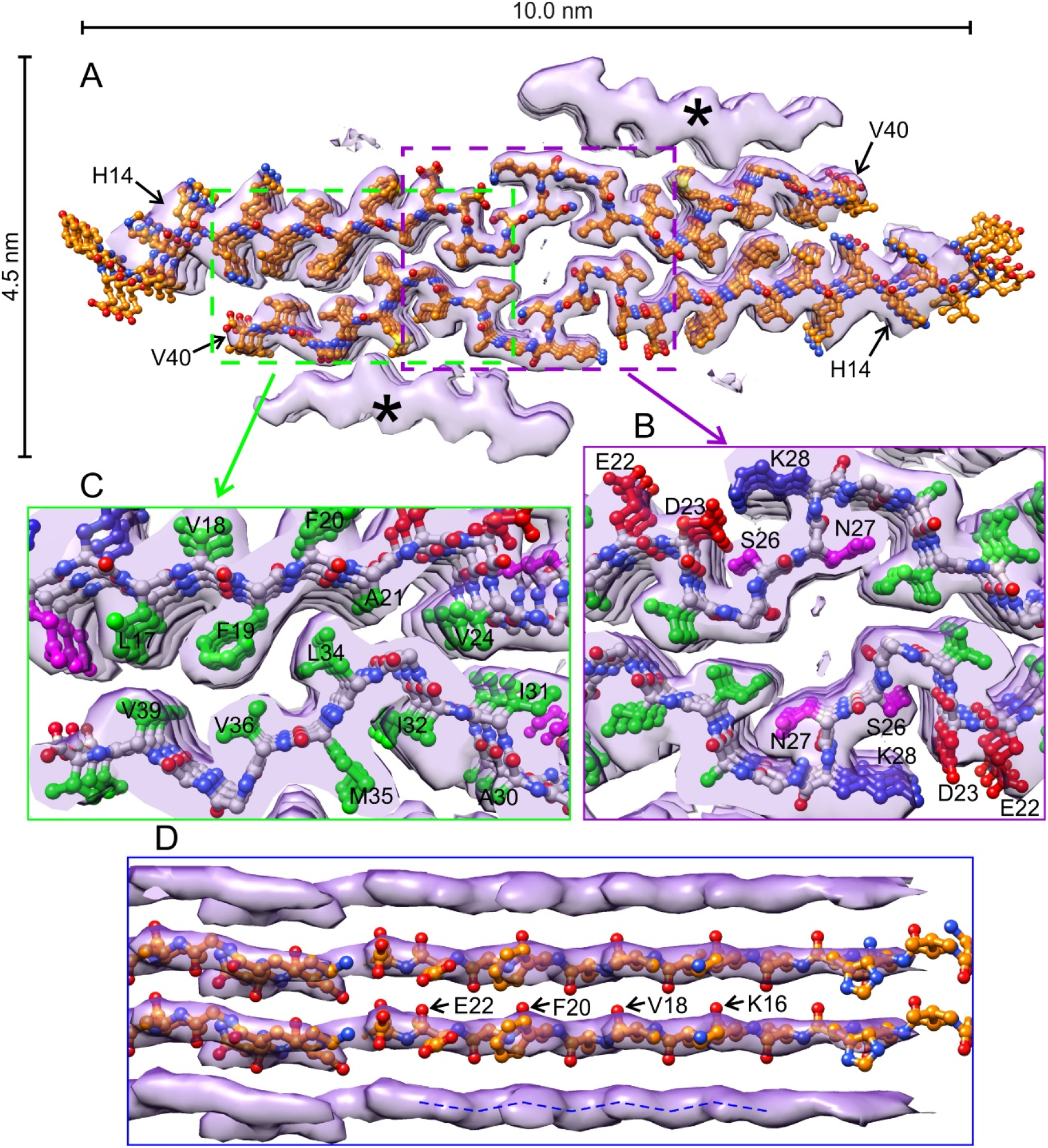
Density map and molecular model for brain-derived Aβ40 fibrils. (A) Cross-sectional view of the full density with three repeats of the molecular model for residues 10-40 in the inner cross-β layers. All carbon atoms are shown in orange. Hydrogen atoms are omitted. The density has near 2_1_ symmetry, with a slight left-handed twist. (B) Expanded view of the central pore, showing D23-K28 salt bridge interactions. Carbon atoms of amino acid sidechains are green (hydrophobic), magenta (polar), blue (positively charged), or red (negatively charged). (C) Expanded view of hydrophobic interactions between the inner layers. (D) Lateral view of the β-sheet formed by residues 14-22. Alignment of backbone density corrugations (dashed blue line) with backbone carbonyl directions favors the left-handed density over its right-handed mirror image.

The density in Fig. 2A consists of four cross-β layers. Amino acid sidechains in the two inner layers are well resolved, allowing a unique fitting of residues 13-40 into the density (*27*). Residues 13-22 form an N-terminal extended segment with one continuous β-strand. Residues 30-40 form a C-terminal extended segment comprised of β-strands in residues 30-32, 34-36, and 38-39, defined by their intermolecular hydrogen bonding patterns. Glycine residues at positions 33 and 37 adopt non-β-strand conformations, disrupting these patterns. All β-strand segments in the inner layers participate in in-register parallel β-sheets. The N-terminal and C-terminal extended segments are separated by an irregular conformation in central residues 23-29, apparently stabilized in part by electrostatic interactions involving oppositely charged sidechains of D23 and K28 (Fig. 2B). Residues 1-12 are not sufficiently ordered to contribute to the density.

Conformations within N-terminal, central, and C-terminal segments in Fig. 2A resemble conformations of the same segments in previously described Aβ fibril structures (*6, 14, 25*), but the overall structural arrangement is qualitatively different. In previous structures, non-β-strand conformations at certain residues allow each Aβ molecule to fold onto itself, resulting in roughly U-shaped (*6, 7, 11, 25*), S-shaped (*12, 13, 15–17*), or C-shaped (*14, 26*) conformations that are stabilized by interactions among hydrophobic sidechains within a single cross-β subunit. In contrast, each Aβ40 molecule in the inner layers in Fig. 2A remains nearly fully extended. Hydrophobic interactions are then exclusively between different cross-β subunits (*i.e.*, different layers of molecules) as shown in Fig. 2C.

The density map in Fig. 2A has a gradual left-handed twist. CryoEM images are invariant to a reflection of the molecular structure, implying that a mirror-image density with a right-handed twist must fit the images equally well. Fig. 2D shows that the left-handed backbone density corrugations match backbone carbonyl directions in our structural model, while corrugations in the mirror-image density (Fig. S3A,B) clash with backbone carbonyl directions. Atomic force microscopy (AFM) images also support a left-handed twist (Fig. S3C).

The inner layers in the density map for our brain-derived Aβ40 fibrils account for 18 kDa/nm of MPL. We propose that the remaining MPL contribution comes from the two outer layers (asterisks in Fig. 2A), implying that each outer layer contributes about 5 kDa/nm to the MPL. The outer layers appear to be cross-β motifs, comprised of β-strands that are 8-9 residues in length, with clear separations along the fibril growth direction. What could the molecular structure in the outer layers be?

One possibility is that the outer layers are cross-β motifs with the usual 4.8 Å intermolecular spacing, but with ~50% occupancy and with only one β-strand segment per molecule. However, the magnitude of the density in the outer layers equals that in the inner layers, ruling out partial occupancy (Fig. S4). A second possibility, that the outer layers are density from residues 1-12 of Aβ40 molecules in the inner layers, is inconsistent with the MPL data. Structural modeling also rules out this possibility, because the outer layers are too far from H13 (Fig. S5).

A third possibility is that the outer layers are cross-β motifs constructed from β-hairpins, resembling structures in fibrils formed by certain designed peptides (*32*). In this context, it is worth noting that Aβ peptides adopt β-hairpin conformations in affibody complexes (*33*), that Aβ peptides constrained to β-hairpin conformations by intramolecular disulfide bridges form oligomers and protofibrils (*34*), that macrocyclic β-hairpin peptides based on Aβ form compact oligomers (*35*), that β-hairpin conformations are sampled in computational studies of Aβ monomers (*36*), and that an early model for Aβ fibril structures was based on β-hairpin conformations (*37*). As depicted in Fig. 3A, a variety of β-hairpin structures are conceivable, all of which would contribute about 4.5 kDa/nm to the MPL.

**Figure 3:**
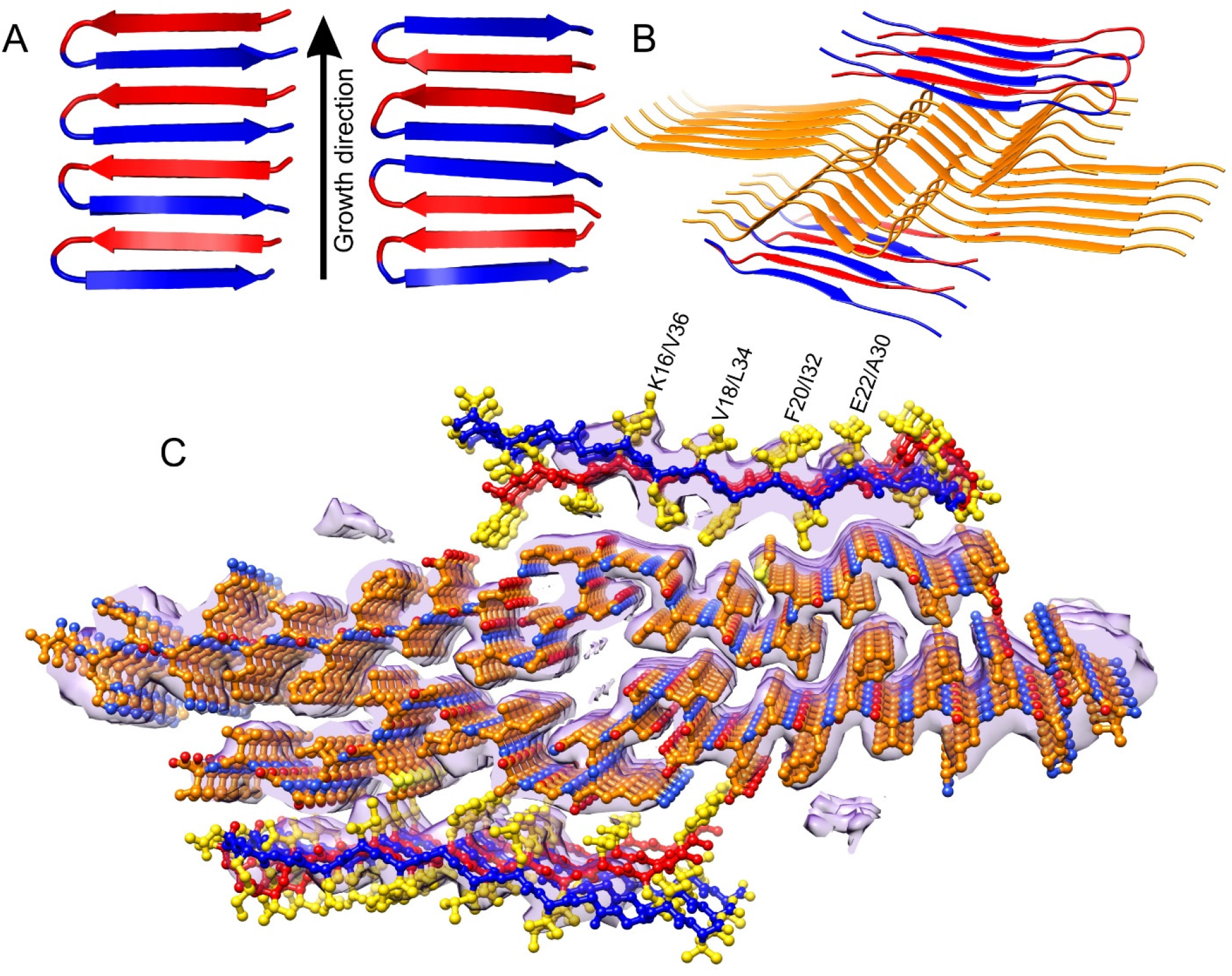
Proposed model for the outer cross-β layers of brain-derived Aβ40 fibrils. (A) Two examples of cross-β motifs comprised of β-hairpins. Any such motif would contribute 4.5 kDa/nm to the experimentally measured MPL value. (B) Ribbon representation of one possibility for the full fibril structure, omitting disordered N-terminal tail segments. Such a structure would have MPL ≈ 27 kDa/nm and would fill the cryoEM density. (C) Atomic representation of the same possible structure. In the outer layers, backbone atoms of residues 13-26 and 27-40 are red and blue, respectively. Sidechain atoms are yellow.

Fig. 3B shows one specific model, in which residues 15-22 and 30-37 form β-strands that participate in both intramolecular and intermolecular backbone hydrogen bonds, and residues 23-29 form turns that lie above or below the C-termini of molecules in the inner layers. Since a fully ordered structure of this type would have a repeat distance of 9.6 Å, densities from the two different β-strand segments of the β-hairpins would be averaged together in Fig. 2A, consistent with the poorly defined sidechains of the outer layers. Attempts to process the cryoEM images with rise values around 9.6 Å did not improve the resolution of these sidechains. However, different fibril segments could contain different β-hairpin-like structures in their outer layers, or different alignments of the β-hairpins between the two outer layers, with only minor effects on the molecular structure of the inner layers. Such disorder in the outer layers would preclude higher resolution.

ssNMR measurements, performed on uniformly ^15^N,^13^C-labeled second-generation fibrils, support the molecular model. From 2D and 3D ssNMR spectra (Figs. 4A, S6, and S7), we obtained ^15^N and ^13^C chemical shift assignments for residues 16-25 and 27-37 (Table S2). Predictions of backbone ϕ and Ψ torsion angles based on these chemical shifts (*38*) are fully consistent with the cryoEM density map (Figs. 4B,C). Torsion angle restraints from ssNMR were included in calculations of a structure bundle by simulated annealing (*27, 39*). The conformation of residues 14-40 in this bundle is quite well defined (Fig. S2C), with root-mean-squared deviation (RMSD) values of 0.37 Å for backbone atoms and 1.37 Å for all atoms.

**Figure 4:**
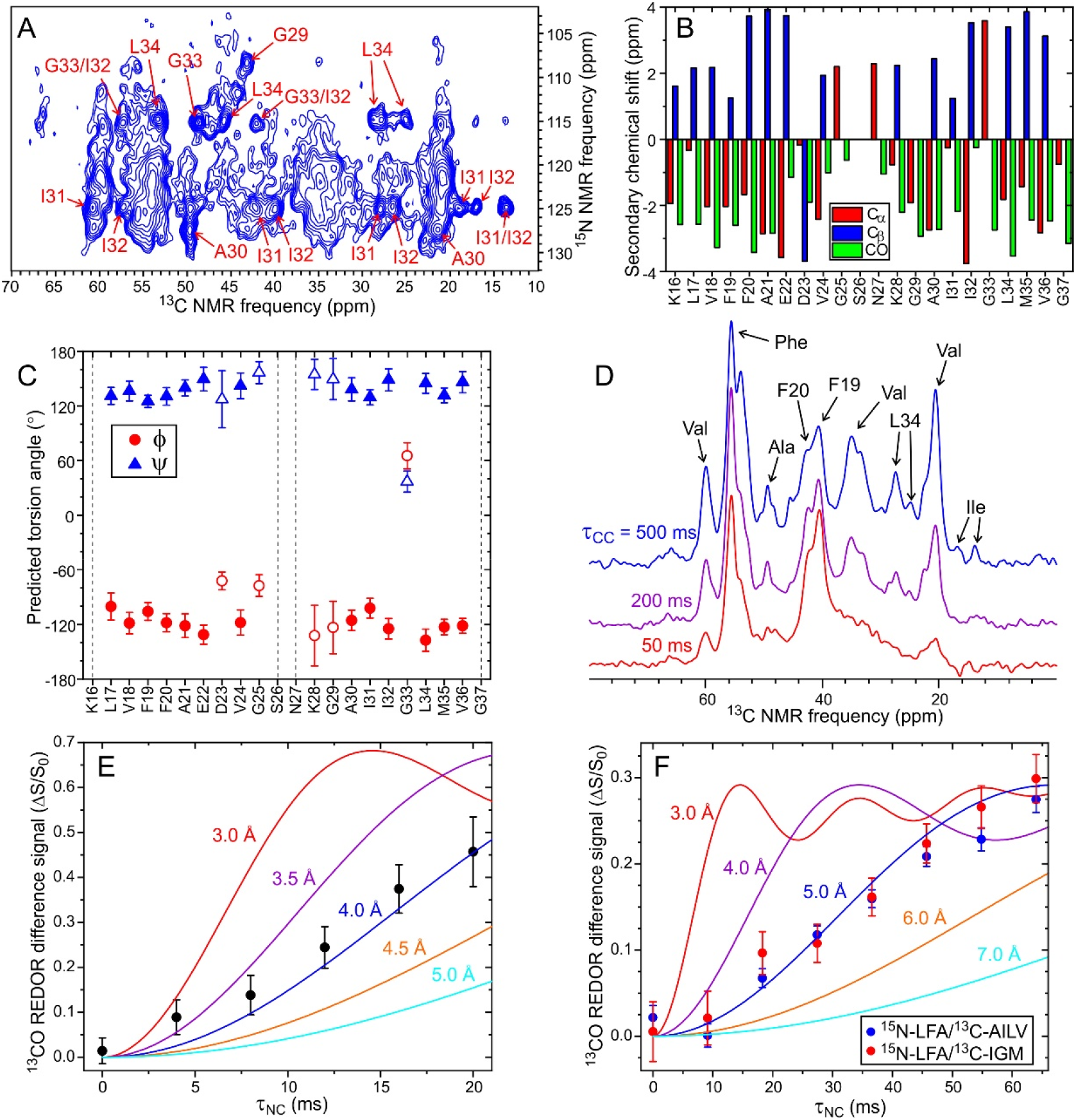
ssNMR data for brain-derived Aβ40 fibrils. (A) 2D NCACX spectrum of second-generation fibrils with uniform ^15^N and ^13^C labeling. Residue-specific assignments for strong, sharp crosspeaks are shown. Contour levels increase by successive factors of 1.25. (B) Secondary ^13^C chemical shifts determined from 2D and 3D spectra. (C) Backbone ϕ and Ψ torsion angle predictions from ^15^N and ^13^C chemical shifts. Predictions classified as “strong” by the TALOS-N program are shown in filled symbols. (D) Aliphatic region of the ^13^C ssNMR spectrum, showing the dependence on the ^13^C-^13^C polarization transfer period τ_CC_ after selective excitation of aromatic ^13^C polarization. (E) Dependence of the fsREDOR difference signal from the sidechain carboxyl site of D23 on the ^15^N-^13^C dipolar dephasing period τ_NC._ Color-coded lines are simulations for dipole-coupled ^15^N/^13^C spin pairs with the indicated interatomic distances. (F) REDOR data for fibrils that were ^15^N-labeled at backbone amide sites of L17, F19, and A21 and ^13^C-labeled at backbone carbonyl sites of A30, I32, L34, and V36 (blue symbols) or I31, G33, and M35 (red symbols). Comparisons with simulations support an average ^15^N-^13^C distance of about 5 Å for one third of the Aβ40 molecules, consistent with a β-hairpin-like structure in the outer layers of the cryoEM density.

2D and 3D ssNMR spectra also contain unassigned signals that may arise from molecules in the outer cross-β layers of the density map (Figs. S6A and S7). Compared to the corresponding spectra of Aβ40 and Aβ42 fibrils in which all molecules are equivalent by symmetry (*13–15, 17, 25*), 2D and 3D ssNMR spectra of the brain-derived Aβ40 fibrils discussed here exhibit relatively broad crosspeaks, despite the fact that the fibrils are morphologically homogeneous in negative-stain TEM and cryoEM images. Greater spectral congestion and broader lines are consistent with the coexistence of structural order in the inner layers with inequivalent molecular conformations and greater disorder in the outer layers.

Inter-residue ^13^C-^13^C polarization transfers, observed in one-dimensional ^13^C spectra after selective excitation of aromatic ^13^C polarization (Figs. 4D and S8) or in 2D ^13^C-^13^C spectra with a 500 ms mixing period (Fig. S6C), indicate contacts between the L34 sidechain and a phenylalanine sidechain (most likely F19), as in Fig. 2C. Measurements of ^15^N-^13^C dipole-dipole couplings with the frequency-selective rotational echo double-resonance (fsREDOR) technique (*40*) indicate a distance of 3.8 ± 0.2 Å between C_γ_ of D23 and N_ε_ of K28 (Fig. 4E), consistent with the D23-K28 salt bridges in Fig. 2B.

Finally, as a test for β-hairpin-like structures in the outer layers of the density map, we prepared second-generation Aβ40 fibrils with ^15^N labels at backbone amide sites of L17, F19, and A21 and ^13^C labels at backbone carbonyl sites of either A30, I32, L34, and V36 or I31, G33, and M35. With these labeling patterns, the shortest ^15^N-^13^C distances within the inner cross-β layers are either 6.5 Å or 8.5 Å (^15^N-A21 to ^13^CO-G33 or^15^N-A21 to ^13^CO-L34 between layers). In contrast, ^15^N-^13^C distances in the 4-6 Å range may occur in a cross-β motif comprised of β-hairpins (Fig. S9). As shown in Fig. 4F, REDOR measurements (*41*) on both selectively labeled fibril samples indicate ^15^N-^13^C couplings that are consistent with an average distance of 5.1 ± 0.3 Å for the carbonyl ^13^C labels in one third of the Aβ40 molecules.

Two other brain-derived Aβ40 fibril structures have been described previously. Using ssNMR, Lu *et al*. developed a structural model for brain-derived fibrils with MPL ≈ 27 kDa/nm, but which were otherwise quite different from the fibrils discussed above (*25*). In particular, the fibrils studied by Lu *et al*. had a rod-like morphology, without width modulation in TEM images, and exhibited a single set of strong, sharp ssNMR signals for residues 1-40, indicating three-fold symmetry about the fibril growth axis (Fig. S10A). The same polymorph was not detected in subsequent measurements by Qiang *et al*. (*24*), demonstrating that this was a rare polymorph, consistent with the atypical clinical history of the AD patient from which it was derived (*25*). Using cryoEM, Kollmer *et al*. developed a structural model for Aβ40 fibrils isolated from meninges (*26*). Unlike the cerebral cortex-derived fibrils discussed above, the fibrils from meninges exhibited a right-handed twist, with a crossover distance of about 40 nm. In the model developed by Kollmer *et al*., residues 1-35 are contained within the cryoEM density and the C-terminus of Aβ40 may also be structurally ordered (Fig. S10B). D23-K28 salt bridges and F19-L34 hydrophobic contacts are absent. Although the density map obtained by Kollmer *et al*. resembles the density map in Fig. 2A, it is not identical. This polymorph may be specific for vascular amyloid.

Since the Aβ40 fibril polymorph discussed above has been shown to be prevalent in the cerebral cortex of most AD patients (*24*), the molecular structure in Fig. 2 can be used to guide the development of amyloid imaging agents with improved specificity and diagnostic utility. The pronounced structural differences among Aβ40 polymorphs may allow the development of structurally-specific aggregation inhibitors (*42*) and antibodies (*43*), targeting polymorphs with the greatest relevance to AD.

## Materials and Methods

### Preparation of Aβ40 fibrils

Aβ40 fibrils for electron microscopy, ssNMR, and AFM were prepared by seeded growth at 24° C in incubation buffer (10 mM sodium phosphate buffer, pH 7.4, 0.01% w/v sodium azide). Second-generation fibrils were prepared from first-generation fibrils described in earlier publications. First-generation “patient 2” fibrils from Lu *et al.* (*25*) were used to prepare uniformly ^15^N,^13^C-labeled second-generation fibrils for ssNMR. First-generation “t-AD3p’” fibrils from Qiang *et al*. (*24*) were used to prepare second-generation fibril samples for cryoEM and AFM and ssNMR. Second-generation t-AD3p’ fibrils were used to prepare third-generation fibrils for MPL measurements. Previously published work shows that Aβ40 fibrils derived from cortical tissue of patient 2 or from t-AD3p’ tissue have indistinguishable 2D ssNMR spectra (*24, 25*), as do first-generation, second-generation, and third-generation fibrils (*28*). Recombinant Aβ40, obtained from Alexotech AB, was used for the uniformly labeled fibrils. Synthetic Aβ40, prepared as described previously (*24, 25, 28*) on Protein Technologies Tribute or Biotage Initiator+ Alstra synthetizers, was used for all other samples.

Tissue samples used to create first-generation fibrils were provided by Dr. Stephen C. Meredith (University of Chicago) and Dr. John Collinge (University College London and Medical Research Council Prion Unit). Production of first-generation fibrils is described in the earlier publications (*24, 25*).

Seeds (*i.e.*, fibril fragments, 50-100 nm in length) were created by sonication of fibrils in incubation buffer (Branson S-259A sonifier with tapered 1/8” microtip horn, lowest power, 10% duty factor, 10-20 min). Lyophilized Aβ40 was dissolved in dimethyl sulfoxide at a typical concentration of 5 mM, then diluted rapidly into incubation buffer containing seeds with a volume sufficient to produce an initial soluble Aβ40 concentration of 50 μM (for electron microscopy and AFM) or 100 μM (for ssNMR). The initial ratio of soluble Aβ40 molecules to Aβ40 molecules in the seeds was between 10:1 and 20:1. Fibril growth proceeded for 18 h before cryoEM grid preparation, 24 h before grid preparation for dark-field TEM, and at least two days before ssNMR or AFM measurements. In all cases, negative-stain TEM was used to confirm the creation of abundant fibrils with lengths exceeding 1 μm and uniform morphologies.

### Negative-stain and dark-field TEM

TEM images were obtained with an FEI Morgagni microscope, operating at 80 keV and equipped with side-mounted Advantage HR and bottom-mounted XR-550B cameras (Advanced Microscopy Techniques). For negative-stain TEM (Figs. 1A and S1A,C), fibril solutions were diluted by factors of 5-10 in deionized water. Aliquots (8-10 μl) were then adsorbed for 1-3 min on glow-discharged TEM grids. Either 300 mesh carbon coated Quantifoil S200/700 grids (Figs. 1A and S1A) or grids consisting of carbon films supported by lacey carbon on 300 mesh copper (Fig. S1C) were used. Grids were blotted, rinsed twice with 8-10 μl of deionized water, then stained with an 8-10 μl aliquot of 2% w/v uranyl acetate for 15-30 s. Grids were then blotted and allowed to dry in air. Images were recorded with the Advantage HR camera.

Dark-field TEM images for MPL measurements (Fig. 1D) were obtained similarly, but using a carbon film with an approximate thickness of 2 nm, coadsorbing 5 μl of diluted fibril solution and 5 μl tobacco mosaic virus (TMV) suspension on the grid for 3 min, rinsing three times with deionized water before final blotting and drying in air, and omitting the uranyl acetate staining step. Dark-field images were recorded with the XR-550B camera, using the tilted-beam conditions described previously (*29*). Care was taken to ensure uniform illumination by the electron beam across the field of view, so that image intensities in different areas of a dark-field image could be treated equally in the MPL analysis.

### Mass-per-length determination

Each MPL value V for the histogram in Fig. 1E was determined from the expression V=(131kDa / nm)×[F−0.5(B_1_ +B_2_)]/ T_ave_, where F is the integrated image intensity within a 100 nm × 40 nm rectangle centered on an Aβ40 fibril segment, B_1_ and B_2_ are integrated background intensities within 100 nm × 40 nm rectangles on either side of the fibril segment, and T_ave_ is the average integrated intensity within 100 nm × 40 nm rectangles centered on TMV rods after subtracting background intensities. Each MPL error value E for the histogram in Fig. S1D was determined from the expression 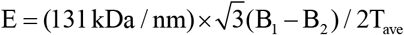, where B_1_ and B_2_ are integrated background intensities within adjacent 100 nm × 40 nm rectangles.

Third-generation t-AD3p’ fibrils were used for MPL measurements because second-generation t-AD3p’ fibrils were found to have too much extraneous material from the original brain extract adhering to their surfaces. This extraneous material was obvious in the dark-field TEM images themselves, resulting in high background intensities and clear variations in intensities along the length of individual fibrils, as well as obvious intensity variations between fibrils. Residual extraneous material associated with third-generation fibrils can account for the difference between the experimental MPL value (28.8 ± 0.4 kDa/nm, Fig. 1E) and the ideal value (27.0 ± 0.5 kDa/nm). Differences in the quantity of extraneous material associated with different fibril segments can also account for the excess width of the MPL histogram in Fig. 1E (13.5 kDa/nm FWHM), relative to the width of the MPL error histogram in Fig. S1D (8.8 kDa/nm FWHM).

It should be noted that second-generation fibrils from patient 2 were found by Lu *et al*. to have MPL = 29.1 ± 1.2 kDa/nm (*25*), indistinguishable from the MPL of third-generation t-AD3p’ fibrils in Fig. 1E. MPL values of approximately 27 kDa/nm have also been measured for certain non-brain-derived Aβ40 fibril polymorphs, using three different electron microscopy techniques that yield dark-field images of unstained samples (*5, 7, 29*). It should also be noted that disordered protein segments (as well as ordered segments) contribute fully to experimentally determined MPL values of amyloid fibrils, as amply demonstrated by previous studies (*29, 44*).

### CryoEM data acquisition

For cryoEM images, 2.5 ul of second-generation Aβ40 fibrils at a concentration of 0.43 mg/ml (100 μM) was applied to a glow-discharged holey carbon grid (Quantifoil R1.2/1.3, 300 mesh gold) for 60 s. The grid was blotted for 6 s, then frozen in cold liquid ethane using a Leica plunger. CryoEM images were acquired with a Titan Krios microscope, operating at 300 keV, using a GIF Quantum K2 camera in super-resolution counting mode and SerialEM software. Each image consists of 50 movie frames with a total exposure time of 10 s, an electron dose of 75 Å^−2^, and a pixel size of 0.54 Å. 1337 images were acquired with defocus values ranging from −0.5 to −3.0 μm.

### CryoEM image processing

CryoEM images were corrected for gain reference and for beam-induced motion and dose-weighted using MOTIONCOR2. Images were binned by a factor of two, yielding a pixel size of 1.08 Å. GCTF was used to estimate the defocus parameters. Fibril segments were picked manually in RELION 3.0. Overlapping particle boxes were extracted using a box length of 400 pixels (432 Å) and 93% overlap (29 Å inter-box distance). Subsequent data processing steps (2D and 3D classification, refinement, and post-processing) were performed with RELION 3.0, a modified version of RELION 3.0-beta, and our own MATLAB programs. Relevant parameters are summarized in Table S1. Regularization parameter values were T = 1 in 2D classification and T = 3 in 3D classification. A full description of 3D density map reconstruction with the modified RELION 3.0-beta software is given in Supporting Text.

In an attempt to identify specific β-hairpin structures in the outer cross-β layers (Fig. 3), we tried several runs with rise values approximately equal to 2 × 4.8 Å = 9.6 Å. The resulting density maps were lower in resolution, without other significant differences. As discussed in the main text, disorder in the outer layers would make it difficult to resolve the β-hairpin structures. Disorder could take the form of coexistence of different β-hairpin motifs (as in Fig. 3A), variations in the identities of β-strand segments or inter-strand hydrogen bonding patterns, variations in the alignment of β-hairpin structures in the two outer layers relative to one another, or variations in the orientations of the β-hairpins relative to the inner layers.

### Initial density model

As the initial model for RELION calculations, we used a lower-resolution density map obtained for *in vitro* Aβ40 fibrils with a similar twisted morphology (*i.e.*, fibrils grown without brain material), low-pass filtered to 10 Å (Fig. S11A). This density map was developed from cryoEM images with a 0.86 Å pixel size, so that its dimensions were effectively expanded by a factor of 1.25 relative to our cryoEM images of brain-derived fibrils.

We also generated an alternative initial model from well-resolved 2D classes of the brain-derived fibrils (Fig. S11C). This was done by aligning the 2D class images manually to form a complete picture of a 2D projection of the fibril. From this picture, approximate values of the rotation angle about the fibril growth direction were assigned to each 2D class, assuming a 180° rotation between crossover points. A MATLAB script (rot_set.m) was then used to insert these rotation angles into the RELION AngleRot variable for boxes that belonged to each well-resolved 2D class. Boxes from poorly resolved 2D classes were not used. We then performed a single 3D classification with unmodified RELION 3.0, using one class, one iteration without image alignment, and a featureless cylinder as the initial model. The resulting density map could be used as the initial model for subsequent calculations.

The two different initial models produced highly similar results in the next stage of calculations (Fig. S11B,D), demonstrating that our final cryoEM density map in Fig. 2 does not depend on details of the initial density model.

### Atomic force microscopy

Tapping-mode AFM images in Fig. S3C were obtained in air, using a Veeco MultiMode instrument and Nanoscope IV controller. A Nanosensors PPP-NCHAuD probe was used, with nominal tip radius of 10 nm, nominal force constant of 42 N/m, and 260 kHz resonant frequency. The scanning rate was 2.8 μm/s. A 10 μl aliquot of second-generation t-AD3p’ Aβ40 fibrils, grown at 50 μM as described above and then diluted 10-fold in deionized water, was adsorbed to freshly cleaved mica for 2 min. After blotting of the mica surface, the surface was rinsed with 10 μl of water and blotted. A 10 μl aliquot of 3% w/v uranyl acetate was then placed on the mica surface for 60 s. The surface was then blotted and allowed to dry in air.

The AFM images reveal that the wider surfaces of these brain-derived fibrils adhere preferentially to the mica surface. Twisting of the fibrils on the surface then occurs primarily within ±20 nm of the crossover points. As a result, height profiles along the length of individual fibrils are strongly peaked at the crossover points, rather than showing the more gradual, sinusoidal modulation that one would expect in the absence of preferentially adhesion by the wider surfaces. In AFM images of fibrils that were adsorbed to mica and dried without uranyl acetate staining, the fibrils were broken at their crossover points, creating a series of flat fibril segments with constant heights of 3-4 nm and lengths of 100-150 nm. Uranyl acetate staining apparently prevents fibril breakage that is driven by interactions with the mica surface, perhaps by reducing the dependence of these interactions on orientation about the fibril growth direction.

### solid state NMR

Unless otherwise stated, ssNMR data were acquired at 14.1 T (599.1 MHz ^1^H NMR frequency, 150.7 ^13^C NMR frequency, 60.7 MHz ^15^N NMR frequency), with a sample temperature of 24 ± 1° C using a Varian InfinityPlus spectrometer console and a 1.8 mm magic-angle spinning (MAS) probe produced by the laboratory of Dr. Ago Samoson (Tallinn University of Technology, Estonia). Data in Figs. 4A,D,E, and S6-S8 were obtained with uniformly ^15^N,^13^C-labeled Aβ40 fibrils, seeded with first-generation fibrils from patient 2 as described above. Data in Fig. 4F were obtained with selectively ^15^N,^13^C-labeled fibrils, seeded with first-generation t-AD3p’ fibrils as described above. ^1^H decoupling fields were 90-110 kHz in all measurements, using two-pulse phase modulation (TPPM) (*45*) during t_1_ and t_2_ evolution periods and during acquisition of ^13^C free-induction decay (FID) signals.

2D and 3D ssNMR spectra of uniformly labeled fibrils were obtained with standard radio-frequency (RF) pulse sequences, using cross-polarization (CP) for ^1^H-^13^C, ^1^H-^15^N, ^13^C-^15^N, and ^15^N-^13^C polarization transfers. Unless otherwise stated, the MAS frequency was 13.6 kHz. RF field amplitudes were typically 68 kHz (^1^H) and 54 kHz (^13^C, ^15^N) for ^1^H-^13^C or ^1^H-^15^N CP, 21 kHz (^15^N) and 35 kHz (^13^C) for ^15^N-^13^CO CP or ^13^CO-^15^N CP, and 9 kHz (^15^N) and 22 kHz (^13^C) for ^15^N-^13^C_α_ CP or ^13^C_α_-^15^N CP. Recycle delays were 1.0 s in 2D and 3D data acquisition.

For 2D NCACX and NCOCX spectra, maximum t_1_ values were 8.0 ms, with 127.2 μs increments and 768 scans per FID. ^13^C-^13^C mixing periods were 100 ms, with approximately 14 kHz ^1^H irradiation for dipolar-assisted rotational resonance (DARR) (*46*) during the mixing period. For the 3D NCACX spectrum, the maximum t_1_ value was 3.60 ms, with 212.0 μs increments, the maximum t_2_ value was 6.57 ms with 212.0 μs increments, and 192 scans were acquired per FID. For the 3D NCOCX spectrum, the maximum t_1_ value was 3.71 ms, with 218.0 μs increments, the maximum t_2_ value was 4.58 ms with 130.8 μs increments, and 208 scans were acquired per FID. For 3D CANCX and CONCX spectra, maximum t_1_ values were 3.10 ms, with 100.0 μs increments, maximum t_2_ values were 4.96 ms with 160.0 μs increments, and 64 (CANCX) or 32 (CONCX) scans were acquired per FID. ^13^C-^13^C mixing periods with DARR were 50 ms in 3D CANCX and 15 ms in 3D CONCX.

The 2D ^13^C-^13^C spectrum in Fig. S6B was acquired at 17.5 T (746.4 MHz ^1^H NMR frequency, 187.7 MHz ^13^C NMR frequency) with a 17.00 kHz MAS frequency. The mixing period consisted of 2.82 ms of finite-pulse radio-frequency-driven recoupling (fpRFDR) (*47, 48*) with 18.00 μs rotor-synchronized ^13^C π pulses. The maximum t_1_ value was 5.1 ms, with t_1_ increments of 20.0 μs and 96 scans per FID. The 2D ^13^C-^13^C spectrum in Fig. S6C was acquired at 14.1 T with a 500 ms DARR mixing period, a maximum t_1_ value of 4.2 ms, t_1_ increments of 21.2 μs, and 192 scans per FID.

2D and 3D data were processed with nmrPipe (*49*), using pure Gaussian apodization functions in all dimension (*i.e.*, no resolution enhancement). Gaussian broadening was 1.1-1.3 ppm in ^15^N dimensions and 0.8-1.0 ppm in ^13^C dimensions (except that it was 0.3 ppm in Fig. S6B and 0.5 ppm in Fig. S6C). 2D and 3D spectra were analyzed and plotted with Sparky (available from https://www.cgl.ucsf.edu/home/sparky/). Chemical shift assignments were made manually.

For data in Figs. 4D and S8, ^13^C polarization of aromatic sidechains was prepared by applying a frequency-selective, Gaussian-shaped π pulse (1.0 ms length) at 128.0 ppm on odd-numbered scans after ^1^H-^13^C CP and a ^13^C “flip-back” π/2 pulse. The MAS frequency was 13.6 kHz. ^13^C signals of odd-numbered scans were subtracted from signals of even-numbered scans. ^13^C-^13^C polarization transfers then occurred for a period τ_CC_, with DARR irradiation, before a second ^13^C π/2 pulse was applied and signals were digitized.

Data in Fig. 4E were obtained with the fsREDOR technique developed by Jaroniec *et al*. (*40*), in which a pair of ^15^N π pulses (12.00 μs) was applied in each MAS rotor period τ_R_ during the ^15^N-^13^C recoupling period τ_NC_ after ^1^H-^13^C CP. Phases of these π pulses followed an XY16 pattern (*50*). A single Gaussian-shaped ^13^C π pulse (600 μs) was applied in the middle of τ_NC_ at 180.4 ppm. A Gaussian-shaped ^15^N π pulse (600 μs) was also applied in the middle of τ_NC_ at 33.6 ppm (*i.e.*, on-resonance with the K28 N_ε_ signal) for acquisition of S_1_ signals. This frequency-selective ^15^N π pulse was omitted for acquisition of S_0_ signals, with no other differences between S_0_ and S_1_. Signals were acquired with ^1^H TPPM decoupling after τ_NC_. RF pulses were actively synchronized with MAS to prevent minor fluctuations of the MAS frequency, which was 8.000 ±±0.003 kHz, from affecting the signals at large values of τ_NC_. The τ_NC_ period (not including Gaussian-shaped π pulse lengths) was set to 32nτ_R_, with n = 0, 1, 2, …, 5. The number of scans per FID was 224(1 + n^2^), with a recycle delay of (1.0 + 0.4n) s. Data are plotted in Fig. 4E as (S_0_ −S_1_)/S_0_ ≡ΔS/S_0_. Error bars are uncertainties calculated from the root-mean-squared (RMS) noise in the S_1_ and S_0_ spectra.

Data in Fig. 4F were obtained with the version of REDOR described by Anderson *et al*. (*41*), in which one ^13^C π pulse (10.00 μs) and one ^15^N π pulse (12.00 μs) was applied in each MAS rotor period during τ_NC_ after ^1^H-^13^C CP. Phases of the ^13^C and ^15^N π pulses followed XY16 patterns. A hard ^13^C π pulse (10.00 μs) was applied in the middle of τ_NC_. A hard ^15^N π pulse (12.00 μs) was also applied in the middle of τ_NC_ for acquisition of S_1_ signals, but not for acquisition of S_0_ signals, with no other differences between S_0_ and S_1_. For sensitivity enhancement, ^13^C signals were acquired with pulsed spin-locking after τ_NC_ as previously described (*51*). RF pulses were actively synchronized with MAS to prevent minor fluctuations of the MAS frequency, which was 7.000 ± 0.003 kHz, from affecting the signals at large values of τ_NC_. The τ_NC_ period was set to 64nτ_R_, with n = 0, 1, 2, …, 8. The number of scans per FID was 256(1 + n^2^), with a recycle delay of (1.0 + 0.5n) s. Error bars in Fig. 4F are uncertainties calculated from the root-mean-squared (RMS) noise in the S_1_ and S_0_ spectra.

PSL removes chemical shift differences among ^13^CO labels in the REDOR measurements on selectively ^13^C-labeled fibrils. Consequently, the observed REDOR signals are the sum of signals from all labeled sites, plus an estimated 11-14% contribution from natural-abundance ^13^C signals (*i.e.*, 0.5 natural-abundance ^13^C nuclei per Aβ40 molecule, on average). Chemical shift differences among labeled carbonyl sites are small and not well resolved, even without PSL, so that REDOR data for individual labeled sites could not be obtained. Moreover, ^13^C chemical shifts for Aβ40 molecules in the outer layers of the cryoEM density map, proposed to contain β-hairpin-like structures, are expected to be different from chemical shifts in the inner layers.

Simulations of ideal REDOR curves for ^15^N-^13^C pairs with a variable internuclear distance were performed with our own programs. In Fig. 4E, the simulated ΔS/ S_0_ values are scaled by a factor of 0.67 to account for the fact that only 2/3 of the Aβ40 molecules may participate in D23-K28 salt bridges. In Fig. 4F, the simulated ΔS/ S_0_ values are scaled by a factor of 0.286 to account for the fact that only 1/3 of the Aβ40 molecules may participate in β-hairpin-like structures and to account for natural-abundance ^13^C signal contributions.

For REDOR measurements in Fig. 4F, selectively labeled fibril samples were cooled to approximately −20° C. Similar measurements above 4° C resulted in smaller ΔS/ S_0_ values, consistent with motional averaging of ^15^N-^13^C dipole-dipole couplings in β-hairpin-like structures at the higher temperatures. These observations suggest that cross-β motifs in the outer layers of the cryoEM density map may be partially dynamically disordered. Dynamic disorder would also result in reduced ssNMR signal amplitudes and broadened crosspeaks in the 2D and 3D spectra of uniformly-labeled fibrils, especially for Aβ40 molecules in the outer layers.

### Molecular model development

An initial model for one Aβ40 molecule in one of the inner cross-β layers of the density map was manually constructed in Coot, using real space refinement. This model included residues 10-40, positioned in the density map in a direction that uniquely optimizes the alignment of large sidechains (*e.g.*, sidechains of K16, L17, F19, F20, K28, I31, I32, and M35) with sidechain density sizes and shapes. At 2.77 Å resolution, the shapes of some of the sidechain densities are quite distinctive, especially the phenylalanine and isoleucine pairs (F19 and F20, I31 and I32), and the pattern of large and small sidechains restricts the alignment. No other alignment of Aβ40 molecules into the inner cross-β layers could be found that was consistent with the pattern of sidechain densities. Six copies of this model were generated and roughly fit into three repeats of the EM density, using manual alignment and the “Fit in Map” function of Chimera (*52*) to rotate and translate each molecule as a rigid body. Xplor-NIH (*39*) was then used to anneal the molecular model, including the probDistPot potential energy function of Xplor-NIH to restrain the model to the density map and the CDIH potential function to restrain backbone torsion angles according to predictions based on ssNMR data (Fig. 4C). As we improved the resolution of the density map, lowest-energy structures from previous Xplor-NIH runs were used as starting structures in subsequent runs. In total, eight rounds of annealing were used, with hydrogen atoms being included only in the final Xplor-NIH calculations.

In the final Xplor-NIH calculations, 60 independent structures were calculated, with annealing from 3000 K to 25 K in 12.5 K decrements, with 5000 steps of torsion angle dynamics at each temperature. The probDistPot potential was applied to heavy atoms of residues 13-40 (excluding oxygen atoms and methyl carbon atoms of V40) with a scale factor of 10.0. The scale factor of the CDIH potential was ramped from 0.002 to 3000 during annealing. Torsion angle ranges were determined by the convertTalos program of Xplor-NIH (version 2.53), using its default parameters (widthCushion = 5°, minWidth = 20°) and were at least 2.2 times the uncertainties reported by TALOS-N. PosDiffPot potentials were used to keep molecular conformations identical within each inner cross-β layer (but not between layers). DistSymmPot potentials were used to enforce translational symmetry within each layer. A PosDiffPot potential was also used to keep backbone heavy atoms of residues 12-40 within 2.5 Å of their initial positions, to prevent molecules from “hopping” from one region of high cryoEM density to another during high-temperature dynamics. The RepelPot potential was used for purely repulsive interactions between atoms at their atomic radii. The TorsionDB potential (scale factor ramped from 0.002 to 2.0), HBDB potential (scale factor 1.0), and standard BOND, ANGL, and IMPR potentials were also used. After annealing, energy minimizations in torsion angle variables and in Cartesian variables were performed.

Twenty-five of the 30 lowest-energy structures (out of 60) exhibited no violations of any Xplor-NIH potentials other than minor violations due to deviations from perfect conformational symmetry among Aβ40 molecules. (The five remaining structures had violations of the torsionDB potential.) A bundle of 10 structures was selected for deposition in the Protein Data Bank (code 6W0O, Fig. S2C), chosen randomly from these 25 structures. This bundle is intended to represent the full range of variations among structural models generated with Xplor-NIH that are fully consistent with the cryoEM density map and ssNMR chemical shifts.

## Acknowledgements

This work was supported by the Intramural Research Program of the National Institute of Diabetes and Digestive and Kidney Diseases, National Institutes of Health. CryoEM measurements were performed at the NIH Multi-Institute CryoEM Facility. Calculations of density maps and molecular models used the computational resources of the NIH HPC Biowulf cluster. We thank Dr. Sjors Scheres for constructing an initial density model used in an early phase of this work. We thank Drs. Jenny Hinshaw and Bertram Canagarajah for their generous assistance and advice.

