## Supporting Information for "Molecular structure of a prevalent amyloid-β fibril polymorph from Alzheimer’s disease brain tissue"

Contents:

Supporting Text

Figures S1-S11

Tables S1 and S2

### Supporting Text

#### Helical Reconstruction with Modified RELION 3.0-beta

To develop a high-resolution cryoEM density map for brain-derived Aβ40 fibrils, we used a modified version of RELION 3.0-beta, supplemented by custom-written MATLAB scripts. All software is available upon request from the authors. The main purpose of these software modifications was to include information about correlations of the orientations of particles (*i.e.*, boxes) about the fibril growth axis when particles come from the same fibril segment. A second purpose was to align the images as much as possible at the 2D classification stage before proceeding to 3D classification. The modified helical reconstruction process consisted of a series of steps, described below.

*Step 1: Initial image processing and particle picking*

Following standard procedures, cryoEM images were corrected for gain reference and beam-induced motion and electron dose-weighted using MOTIONCOR2. The images were binned from their original pixel size of 0.54 Å to a final pixel size of 1.08 Å. GCTF was used to estimate the defocus parameters. Fibril segments were picked manually in unmodified RELION 3.0, and 400 pixel X 400 pixel boxes (particles) were extracted from the fibril segments with 93% overlap.

*Step 2: 2D classification and particle alignment*

Fifteen rounds of 2D classification were performed with unmodified RELION 3.0, using a regularization value T = 1 and 70-100 classes. Boxes were removed either as entire poorly-resolved 2D classes or individually after sorting the boxes by RELION Z-score.

After the first round of 2D classification and after two later rounds, 2D class images were examined in ImageJ (*53*). For each well-resolved 2D class image, the angle α between the x-axis of the image and the fibril growth axis and the displacement δ of the fibril in the y-axis direction from the center of the image were measured. Ideally, if all fibril segments were perfectly aligned and centered in their boxes, both quantities should be zero. To correct for non-zero values, two MATLAB scripts (psi_tweak.m and shift_y.m) were used to adjust the RELION orientation and translation variables for each box according to its 2D class, as follows:

$$\begin{aligned} &\text{AnglePsi} \rightarrow \text{AnglePsi} + \alpha \\ &\text{AnglePsiPrior} \rightarrow \text{AnglePsiPrior} + \alpha \qquad (1) \\ &\text{OriginX} \rightarrow \text{OriginX} + \delta\sin(\text{AnglePsi}) \\ &\text{OriginY} \rightarrow \text{OriginY} + \delta\cos(\text{AnglePsi}) \end{aligned}$$

The new values of these variables were stored in an updated "particles.star" file. Note that AnglePsi is the particle orientation angle about the z-axis, *i.e.*, perpendicular to the plane of the image. OriginX and OriginY are particle displacements in the xy plane. (For simplicity, we omit the prefix "_rln" from RELION variables.)

In addition, based on the asymmetry of the fibrils in the x-axis direction about their crossover points, 2D classes were judged by inspection to contain boxes in which fibrils ran in "forward", "backward", or "ambiguous" directions. For each picked fibril segment (identified by the MicrographName and HelicalTubeID values of its boxes), a MATLAB script (modify_psi_nopitch.m) was used to create two sums, F and B. F is the sum of the MaxValueProbDistribution values of boxes from the fibril segment that belong to "forward" classes. B is the same sum from boxes that belong to "backward" classes. If B > F, the script added or subtracted 180° from AnglePsi and AnglePsiPrior values of all boxes in the fibril segment. If all boxes in the fibril segment were assigned to ambiguous 2D classes, the entire segment was discarded. The motivation for these manipulations was to develop and enforce a consensus regarding the most likely direction of the fibril segment from which the individual boxes were extracted. The directionality of fibril segments is not apparent in each cryoEM image but can potentially be revealed by 2D classification.

The same MATLAB script also set initial values for the variable AnglePsiFlipRatio for all boxes in each fibril segment, equal to the greater of $0.5-0.25|B-F|$ or 0.2. The initial AnglePsiFlipRatio value was always between 0.2 and 0.5. This value determines how RELION weights the probabilities of AnglePsi values that differ by 180°.

*Step 3: Initial 3D classification*

As an initial 3D density model, we used a lower-resolution density map from a study of Aβ40 fibrils prepared *in vitro* (*i.e.*, not derived from brain tissue). 2D classes for the *in vitro* fibrils were similar to 2D classes for the brain-derived fibrils. This initial model was low-pass filtered at 10 Å. Unmodified RELION 3.0 was then used for a first 3D classification (K = 1, T = 3), so that each box would get an initial value for the angle AngleRot of its maximum probability alignment. AngleRot represents the angle of rotation about the fibril growth direction.

*Step 4: Adjustment of rotation angles within fibril segments by local averaging*

Because of the helical symmetry, the rotation angle $\rho(\Delta)$ (in degrees) of one box centered at a distance Δ along the growth axis of the fibril from a second box with rotation angle $\rho_0$ should ideally be $\rho(\Delta)=\rho_0+180(\Delta/p)$, where p is the fibril pitch defined as the minimum distance between points whose rotation angles differ by 180°. This is usually the crossover distance, but with a sign that indicates the direction of the fibril ("forward" or "backward" as discussed above).

Using MATLAB scripts (2d_class_rot.m and hist_av.m), we created histograms of the AngleRot values of boxes in each good 2D class. Due to the approximate two-fold symmetry of the fibrils, there were typically two peaks in the histogram, separated by 180°. For each 2D class, we recorded the average AngleRot value from each of the two peaks, as well as a weighting factor that was 0.1 for 2D classes without well-defined histogram peaks and 1.0 otherwise.

Next, we assigned tentative rotation angles to each box. Certain 2D class images did not have full mirror symmetry about their midlines (*i.e.*, about the x-axis) and occurred as mirror-image pairs. For a few of these classes, overlapping regions could be identified, so the 2D classes could be linked together into two mirror image groups. Boxes from one group were assigned the AngleRot value from one of its histogram peaks, while boxes from the mirror-image group were assigned an AngleRot value that differed by 180°. Boxes from other 2D classes were assigned AngleRot values from the histogram peak that was most consistent with the other boxes extracted from the same fibril segment, using a MATLAB script (rot_assign.m). Here, "consistent" means that the equation $\rho(\Delta)=\rho_0+180(\Delta/p)$ is satisfied as nearly as possible. Values of Δ were determined from the variable HelicalTrackLength. If the fibril segment did not contain any boxes from the 2D classes that were assigned a mirror-image group, the choice of which AngleRot histogram peak value was chosen randomly for the first box and the rest of the AngleRot values chosen for consistency.

The same MATLAB script also set the translation alignment along the fibril growth direction of all boxes to zero.

$$\begin{aligned} &Q = \text{OriginX} \sin(\text{AnglePsi}) + \text{OriginY} \cos(\text{AnglePsi}) \\ &\text{OriginX} \rightarrow Q\sin(\text{AnglePsi}) \\ &\text{OriginY} \rightarrow Q\cos(\text{AnglePsi}) \end{aligned} \quad (2)$$

This was done to avoid wasted image data that can result from boxes being shifted outside of the bounds of the 3D model during RELION calculations. When OriginX and OriginY values were changed, corrections to the assigned AngleRot values were also made as required by the helical symmetry.

Finally, local averaging of the tentatively assigned AngleRot values within each fibril segment was performed. Using another MATLAB script (smooth_rot.m), a new rotation angle was determined for each box by a weighted average of the rotation angles of other boxes from the same segment, using the following equations:

$$\begin{aligned} &C = \sum_{\Delta} w_{\Delta} \cos[\rho(\Delta) - 180(\Delta / p)] \\ &S = \sum_{\Delta} w_{\Delta} \sin[\rho(\Delta) - 180(\Delta / p)] \\ &\rho_{0,new} = \frac{S}{|S|} \cos^{-1}(C / \sqrt{C^2 + S^2}) \\ &w_{\Delta} = a_{\Delta} \exp(-\Delta^2 / 2\sigma^2) \end{aligned} \quad (3)$$

In Eqs. (3), $\sum_{\Delta}$ represents a sum over boxes in the same fibril segment, displaced by Δ from the box under consideration, and $a_{\Delta}$ is the weighting factor determined from AngleRot histograms as explained above. σ was chosen to be 100 pixels to provide averaging primarily among nearby boxes. Angular averaging is done by averaging the cosine and sine components to avoid problems caused by the discontinuity between 180° and -180°. Local averaging described by Eqs. (3) was repeated until no rotation angle changed by more than 0.1°. Final values of $\rho_{0,new}$ were then used to update the AngleRot and AngleRotPrior values of each box, prior to further 3D classification with the modified version of RELION 3.0-beta described below.

*Step 5: Further 3D classification with modified RELION software*

For further 3D classification, we modified RELION 3.0-beta to include local averaging of the rotation angle among boxes that were extracted from the same fibril segment, as in step 4.

RELION already has the option for local averaging of AnglePsi values. The implementation of averaging of AngleRot values is similar. Modifications were made primarily in RELION's UpdatePriorsForOneHelicalTube and HealpixSampling functions. With these modifications, AngleRotPrior values were locally averaged after each iteration of model optimization within RELION 3.0-beta, using the following equations:

$$\begin{aligned} C &= \sum_{\Delta} w_{\Delta} \cos[\rho(\Delta) - 180[(\Delta + \Delta_{trans}) / p]] \\ S &= \sum_{\Delta} w_{\Delta} \sin[\rho(\Delta) - 180[(\Delta + \Delta_{trans}) / p]] \\ \rho_{0,new} &= \frac{S}{|S|} \cos^{-1}(C / \sqrt{C^2 + S^2}) \\ w_{\Delta} &= P_{max}(\Delta) \exp(-\Delta^2 / 2\sigma^2) \end{aligned} \quad (4)$$

Eqs. (4) are identical to Eqs. (3), except that the difference in most probable translations $\Delta_{trans}$ is added to the initial displacement between boxes Δ, the pitch value p (in units of pixels per 180° helical rotation) is contained in RELION's HelicalTubePitch variable, $P_{max}(\Delta)$ is the maximum value of the alignment probability distribution of each box (stored in RELION variable MaxValueProbDistribution), and σ is calculated as RELION's "range factor of local averaging", which we set to 0.4 in units of the box length. When $|\Delta| > 3\sigma$, $w_{\Delta}$ is set to zero.

Because of the near two-fold screw symmetry of the fibrils, the rotation angle search was designed to cover both the region defined by AngleRotPrior ± "local angular search range," and the region 180° away. This was done in a similar way as the AnglePsi search in unmodified RELION. In the modified RELION 3.0-beta, a new variable AngleRotFlipRatio is used in the same way as the variable AnglePsiFlipRatio to control the relative probability of the two angular search regions. If a majority of the boxes in a fibril segment have their most probable AngleRot values near AngleRotPrior ± 180°, then AngleRotFlipRatio > 0.5. The modified RELION program then adds or subtracts 180° from the AngleRotPrior values of all boxes in the segment and replaces AngleRotFlipRatio with the quantity 1.0 - AngleRotFlipRatio. If AnglePsiFlipRatio is greater than 0.5, both the AnglePsiPrior value and the AngleRotPrior value are changed by ±180°.

3D classification was done with the modified RELION 3.0-beta software, first using one 3D class (K = 1, T = 3). The density map from this run was used as the initial model for a subsequent run with three 3D classes (K = 3, T = 3). The best-resolved 3D class, which included 43% of the boxes (104,672 boxes) from 2D classification, was used for further processing.

Masks were used in 3D classification (after the first one), 3D auto-refinement, and post-processing, in addition to setting the "outer tube diameter" parameter to 145 Å. Because the 3D model is generated in a 432 Å cube, while the widest cross-sectional dimension of the fibrils is about 120 Å, large regions of the 3D model cube contain only noise and can be excluded from the analysis by setting the tube diameter and using a mask. Our masks were generated in RELION from the density of the previous step with a 15 Å low-pass filter, extending the binary map by 8 pixels, and adding a soft edge of 8 pixels. The masks were manually inspected in Chimera to ensure that the masks were not excluding any pixels in the central region of the 3D model.

*Step 6: Additional processing*

Using the modified version of RELION 3.0-beta, 3D auto-refinement was done with only helical symmetry (using the previously generated density map as the initial model, filtered to 10 Å). The resolution at this stage was 3.25 Å after post-processing, with -0.69° helical twist and 4.88 Å rise. CTF refinement was then run to fit the per-particle defocus and the beam tilt estimation. Astigmatism was not adjusted, either at the per-particle or per-micrograph level, because testing showed that the calculated astigmatism adjustments would be unreasonably large. After calculation of the CTF refinements, a MATLAB script (compare_ctf.m) was used to discard any boxes whose defocus adjustment was greater than 600 Å. About 1% of the boxes were discarded, leaving 103,610 boxes.

The motion correction adjustment of the images (particle polishing) was first trained with 5500 boxes, then those parameters were used for the entire dataset. At this stage, we repeated 3D auto-refinement and post-processing with only helical symmetry. The resolution improved to 2.98 Å. We then repeated the entire cycle of CTF refinement, motion correction, 3D auto-refinement, and post-processing. This did not make any significant difference, with a final reported resolution of 2.96 Å, with -0.71° twist and 4.89 Å rise. At this point, 103,462 boxes remained.

For the final density map with $2_1$ symmetry (Figs. 2 and S2), we low-pass filtered the final result from processing with only helical symmetry to 15 Å, then ran 3D auto-refinement and post-processing with $2_1$ symmetry. We used the resulting density, low-pass filtered to 6 Å, as the initial model for 3D classification (K = 1, T = 3), then 3D auto-refinement (also using a 6 Å low-pass filter) and post-processing. The final resolution was 2.77 Å, with -180.34° twist and 2.45 Å rise (corresponding to -0.68° twist and 4.90 Å rise without symmetry).

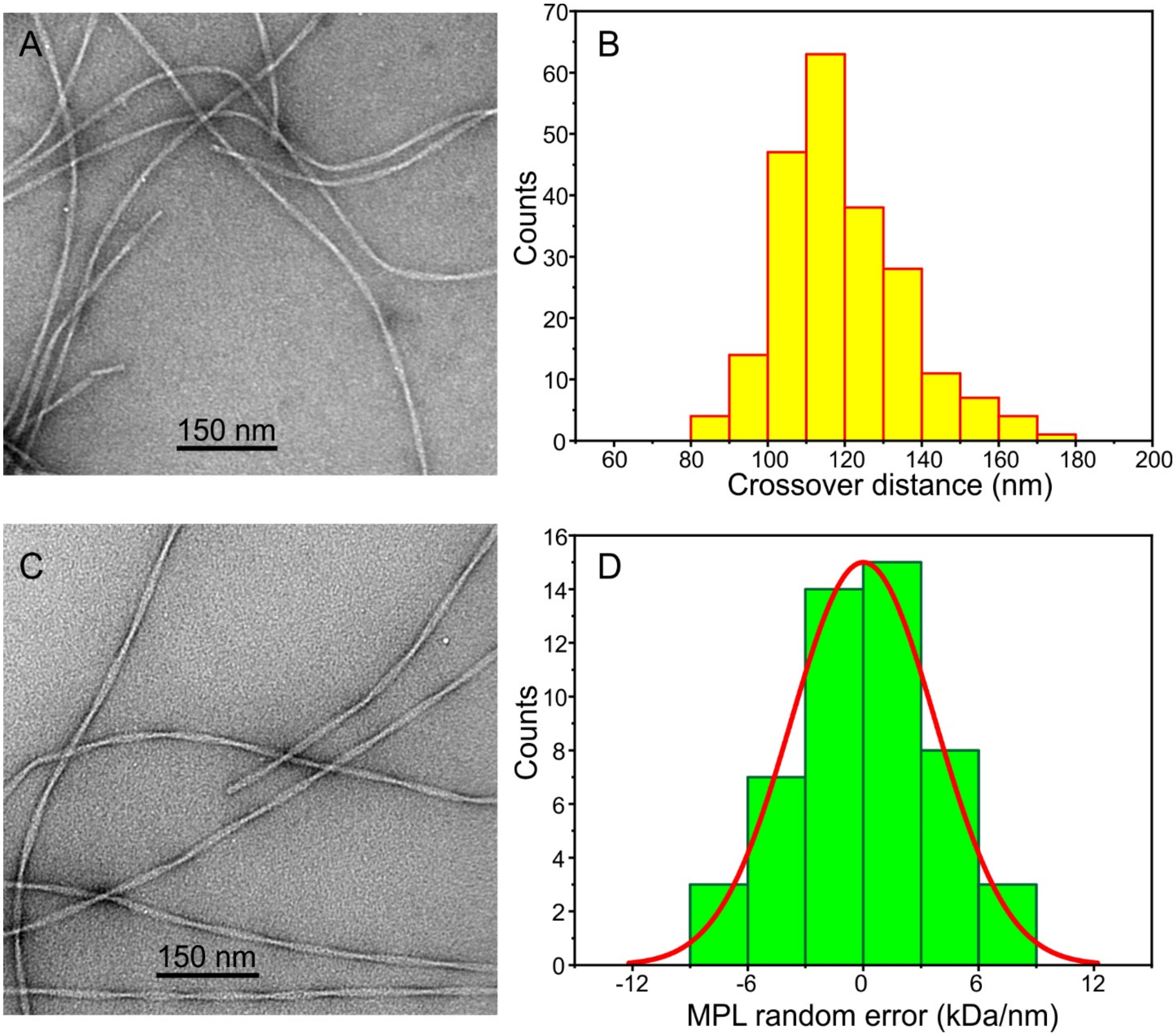


Figure S1: (A) Additional negative-stain TEM image of second-generation Aβ40 fibrils used for cryoEM measurements. (B) Histogram of distances between crossover points for fibrils in cryoEM images. (C) Negative-stain TEM image of third-generation Aβ40 fibrils used for MPL measurements. (D) Histogram of errors in MPL measurements due to random fluctuations in background intensity in dark-field TEM images. Each error value was determined from the difference in integrated image intensity for a pair of adjacent 100 nm × 40 nm rectangular regions, in image areas that were devoid of fibrils. A Gaussian fit centered at 0.0 kDa/nm with 8.8 kDa/nm FWHM is shown in red.

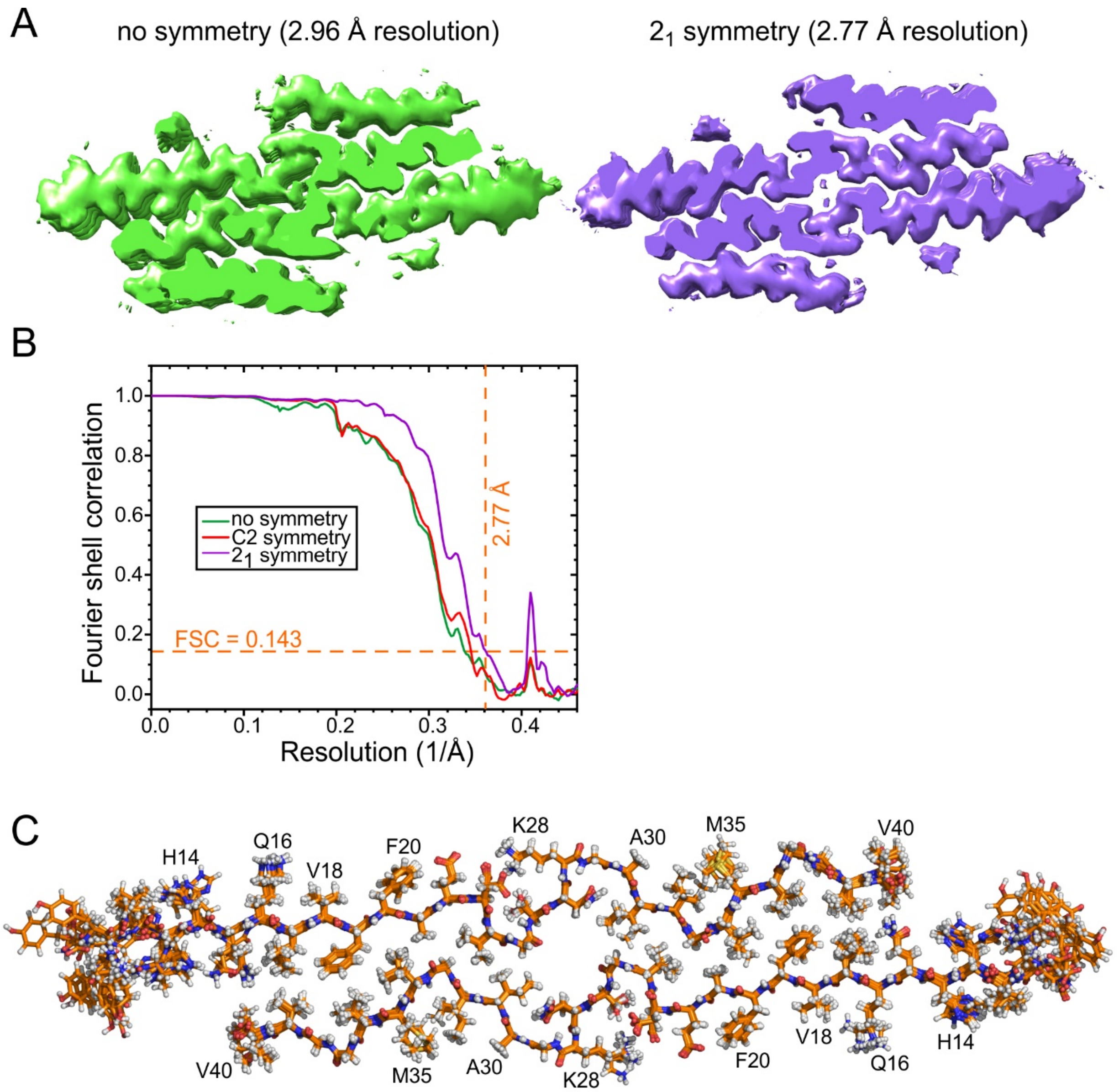


Figure S2: (A) Comparison of cryoEM densities from RELION calculations without rotational or screw symmetry about the fibril growth direction (green, helical rise = 4.89 Å, twist = -0.71°) and with near $2_1$ screw symmetry (purple, helical rise = 2.45 Å, twist = -180.34°). (B) FSC curves for densities calculated without additional symmetry, with near $C_2$ rotational symmetry (helical rise = 4.90 Å, twist = -0.69°), and with near $2_1$ symmetry. Orange dashed lines indicate "gold standard" resolution of 2.77 Å for the density calculated with $2_1$ symmetry. (C) Superposition of molecular models from ten independent simulated annealing calculations with Xplor-NIH, using the density map of the two inner cross-β layers to restrain the backbone and sidechain conformations of residues 13-40 (except $C_\gamma$ of V40). Predictions for backbone $\phi$ and $\psi$ torsion angles of residues 17-22, 24, 30-32, and 34-36 from TALOS-N, based on $^{15}N$ and $^{13}C$ chemical shift assignments from 2D and 3D ssNMR spectra, were used as additional conformational restraints. Atomic coordinates for this structure bundle are available in the Protein Data Bank (code 6W0O).

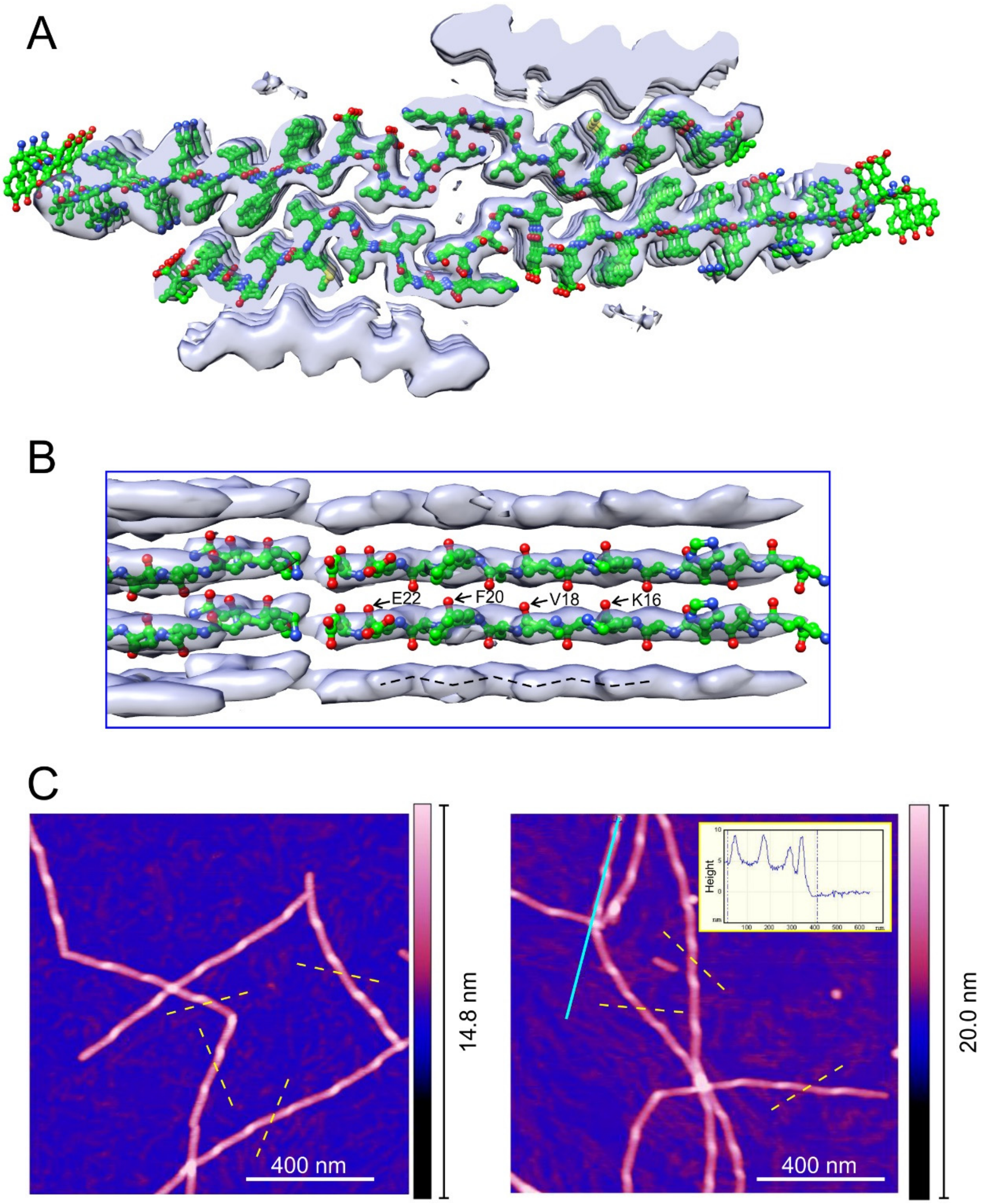


Figure S3: (A) Mirror-image cryoEM density, with a slight right-handed twist, and molecular model for residues 10-40 of Aβ40 in the inner cross-β layers. This molecular model was calculated as in Figs. 2A and S2C, but using the mirror-image density. (B) Lateral view of the in-register, parallel β-sheet formed by residues 14-22. With the mirror-image density, backbone carbonyl directions do not align correctly with corrugations in backbone density. (C) AFM height images of second-generation Aβ40 fibrils. Asymmetries in height profiles around crossover points align with the dashed yellow lines, supporting a left-handed twist. Inset on the right shows a height section along the cyan line.

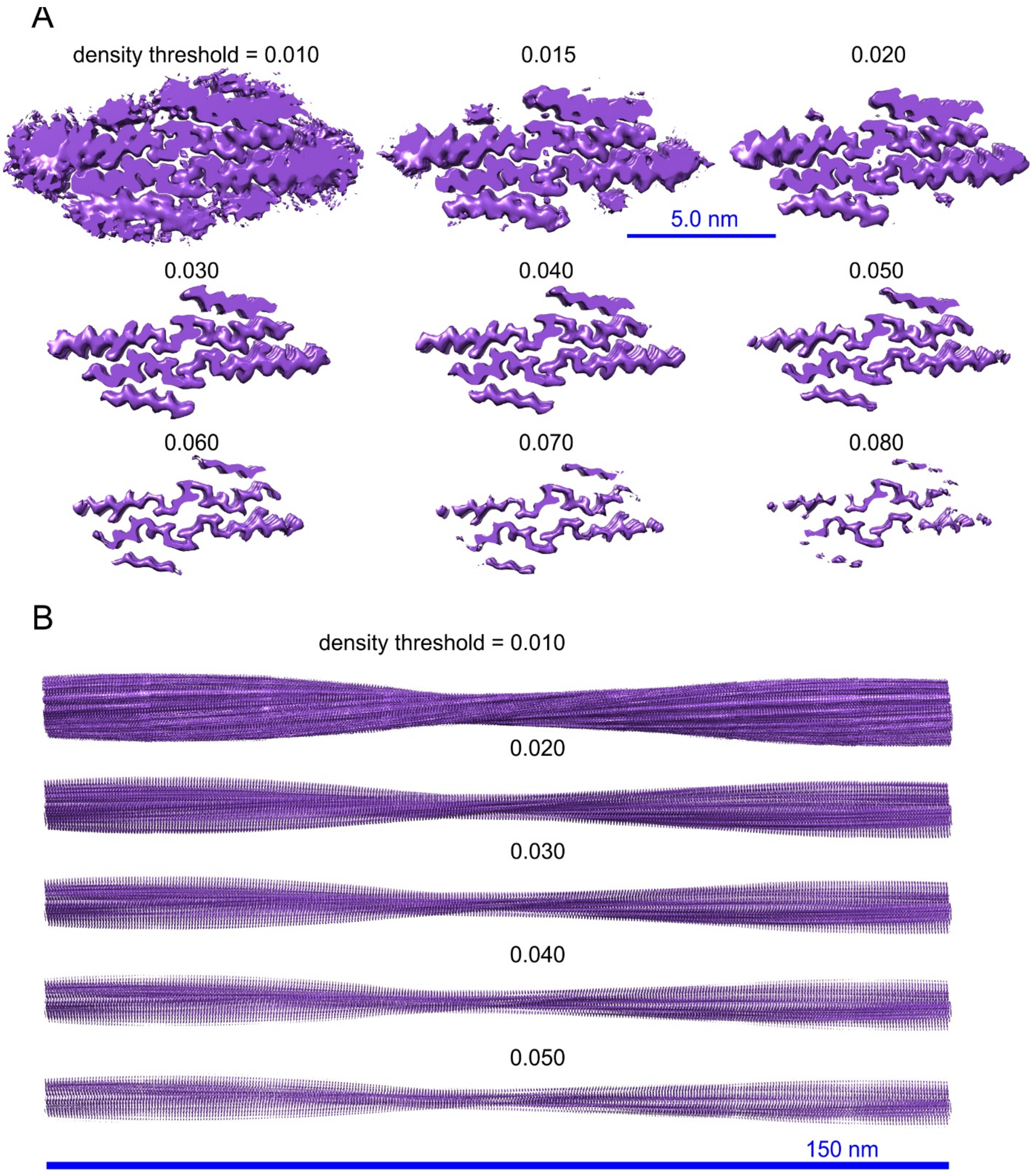


Figure S4: (A) CryoEM density map for brain-derived Aβ40 fibrils represented by surfaces at the indicated threshold values. A cross-sectional slice of the density is shown. Threshold values are on an arbitrary scale, as reported by the Chimera visualization software used to generate these images. (B) Side view of a long fibril created from the cryoEM density map.

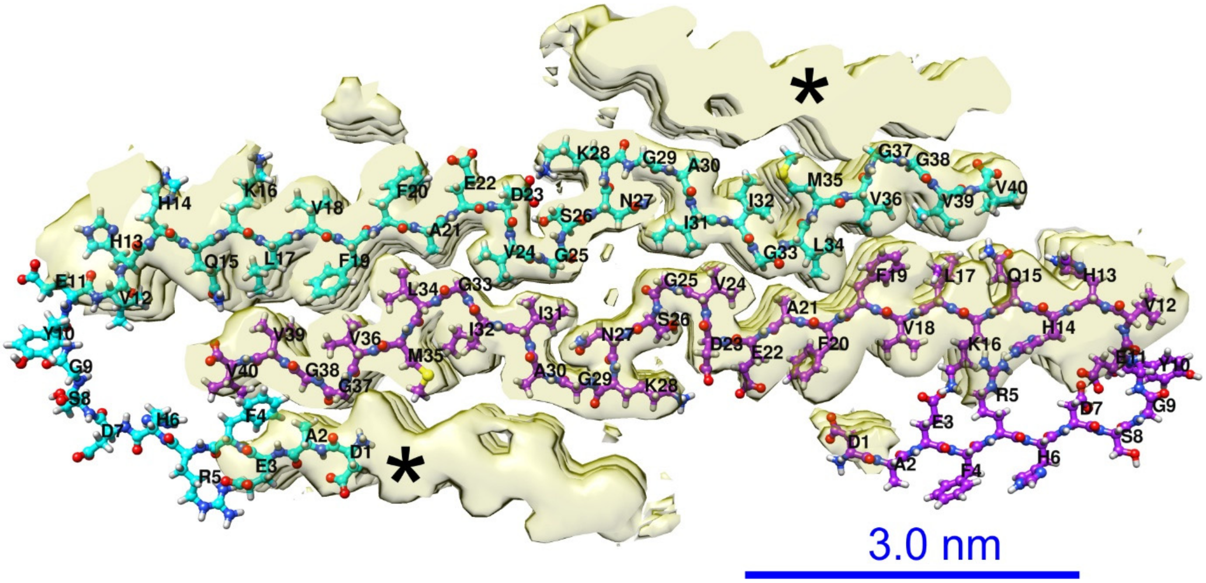


Figure S5: Attempt to fill the outer cross-β layers of the cryoEM density, indicated by asterisks, with N-terminal tails of Aβ40 molecules from the inner layers. This representation shows that the N-terminal tails (residues 1-12) do not have sufficient length to reach the outer layers in either direction.

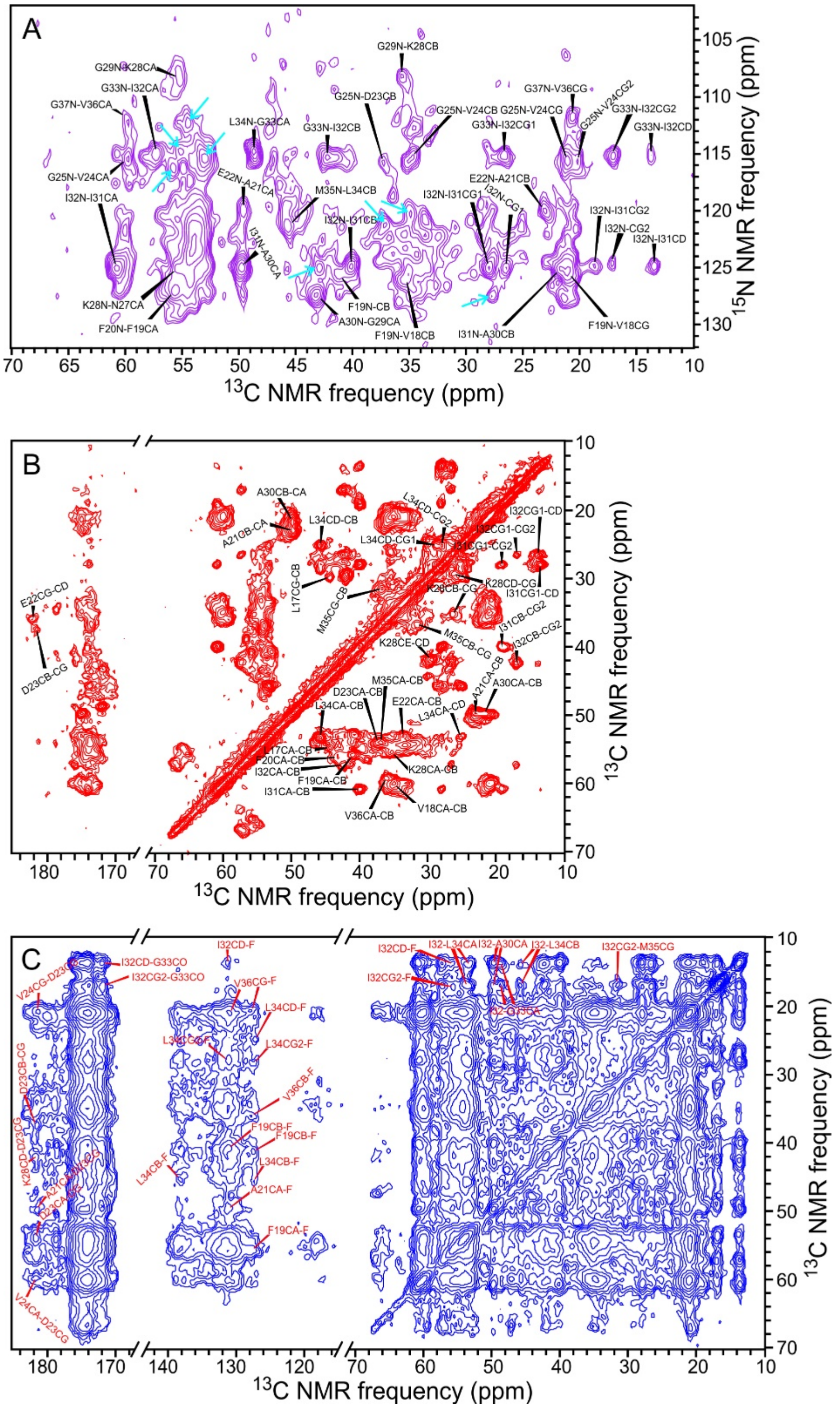


Figure S6: (A) 2D NCOCX ssNMR spectrum of uniformly $^{15}$N,$^{13}$C-labeled second-generation brain-derived Aβ40 fibrils, with assignments of a subset of the crosspeak signals. Cyan arrows indicate examples of crosspeaks that can not be assigned to residues 16-37 of Aβ40 molecules in the inner cross-β layers. Contour levels increase by successive factors of 1.25. (B) 2D $^{13}$C-$^{13}$C spectrum of the same fibrils, acquired with a 2.82 ms fpRFDR mixing period. (C) 2D $^{13}$C-$^{13}$C spectrum acquired with a 500 ms DARR mixing period. Assignments of inter-residue crosspeaks are shown, with assignments to "F" being either F19 or F20.

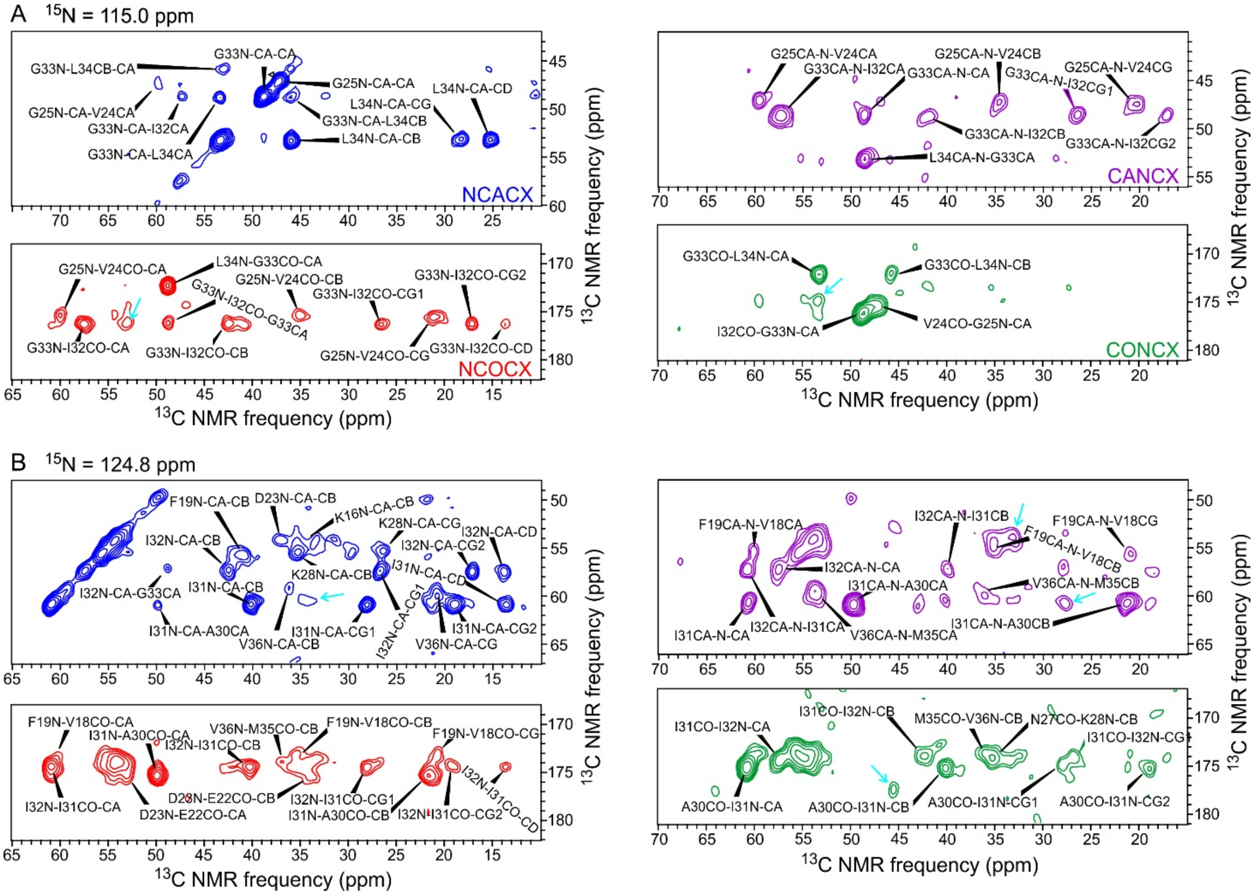


Figure S7: 2D planes from 3D NCACX (blue), NCOCX (red), CANCX (purple), and CONCX (green) ssNMR spectra of uniformly $^{15}$N,$^{13}$C-labeled Aβ40 fibrils. $^{13}$C-$^{13}$C planes are shown at $^{15}$N chemical shifts of 115.0 ppm (A) and 124.8 ppm (B). Cyan arrows indicate examples of crosspeaks that can not be assigned to residues 16-37 of Aβ40 molecules in the inner cross-β layers. Contour levels increase by successive factors of 1.40.

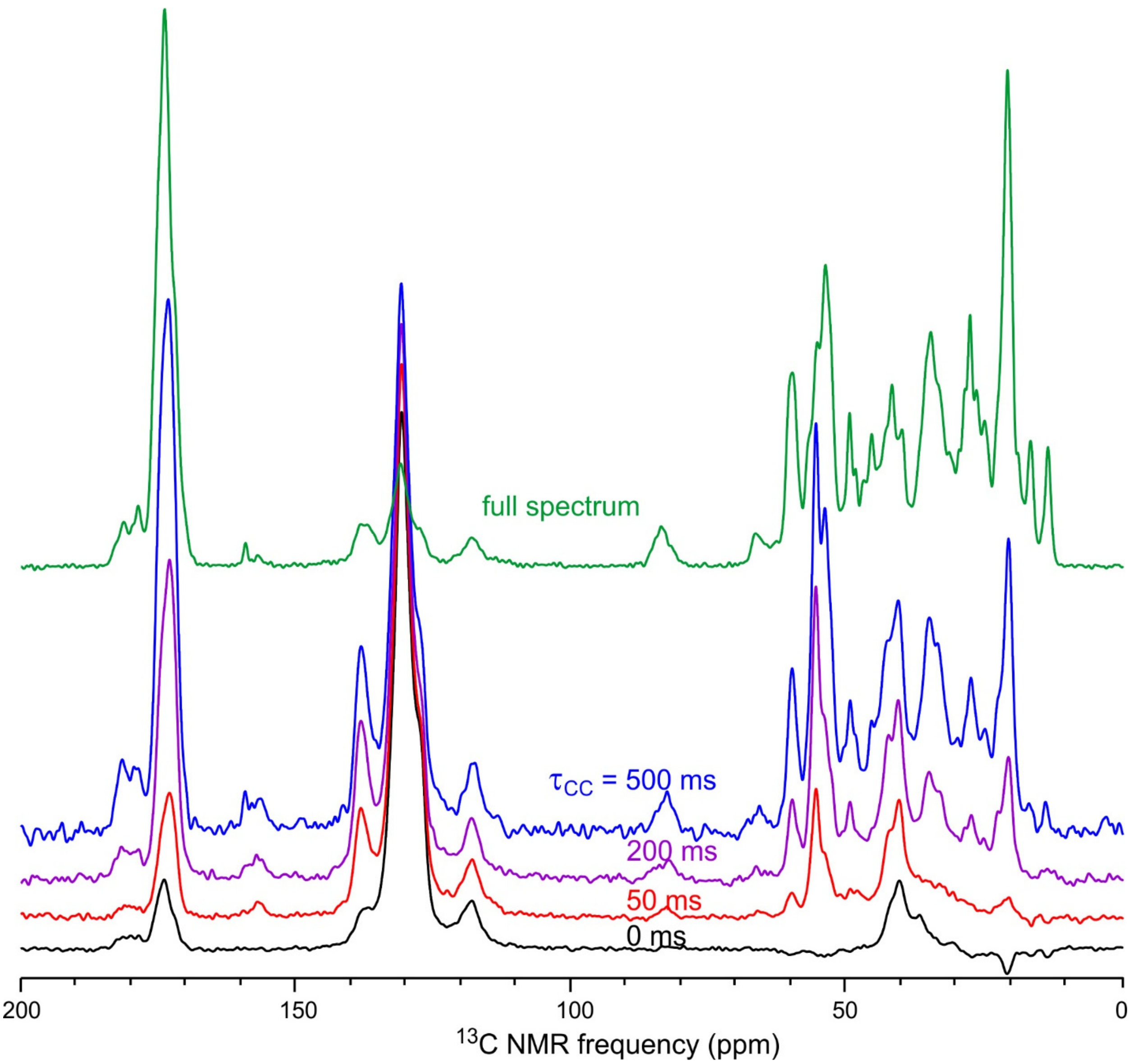


Figure S8: $^{13}$C ssNMR spectra of uniformly $^{15}$N,$^{13}$C-labeled Aβ40 fibrils with the indicated $^{13}$C-$^{13}$C polarization transfer periods $\tau_{CC}$ after selective excitation of aromatic $^{13}$C polarization. The full spectrum, without selective excitation, is shown in green.

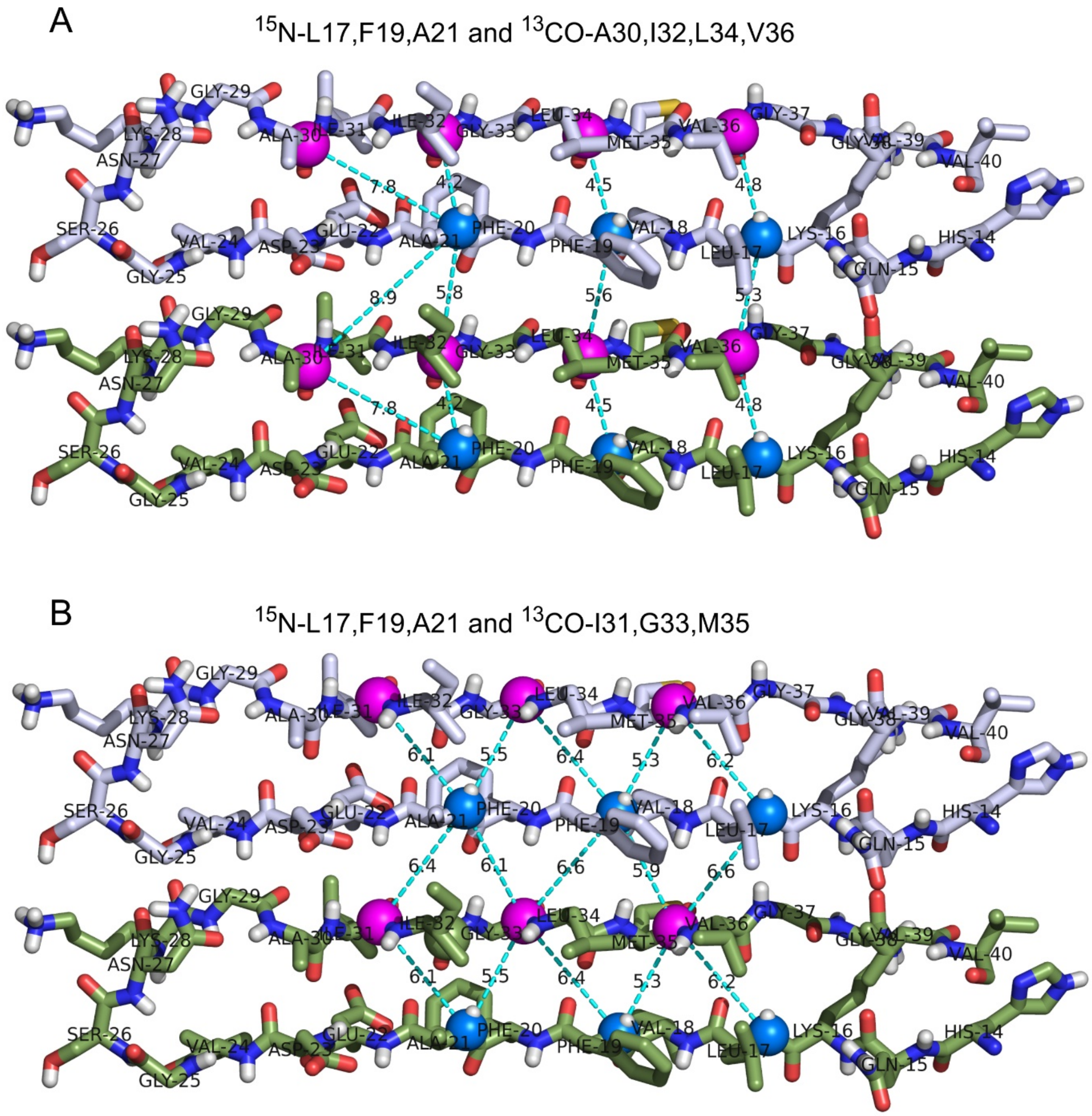


Figure S9: Possible β-hairpin structures for residues 14-40 of Aβ40, showing intramolecular and intermolecular distances between $^{15}$N labels (blue spheres) and carbonyl $^{13}$C labels (magenta spheres) with $^{15}$N labels at L17, F19, and A21 and $^{13}$C labels at A30, I32, L34, and V36 (A) or I31, G33, and M35 (B). Distances are shown in angstrom units.

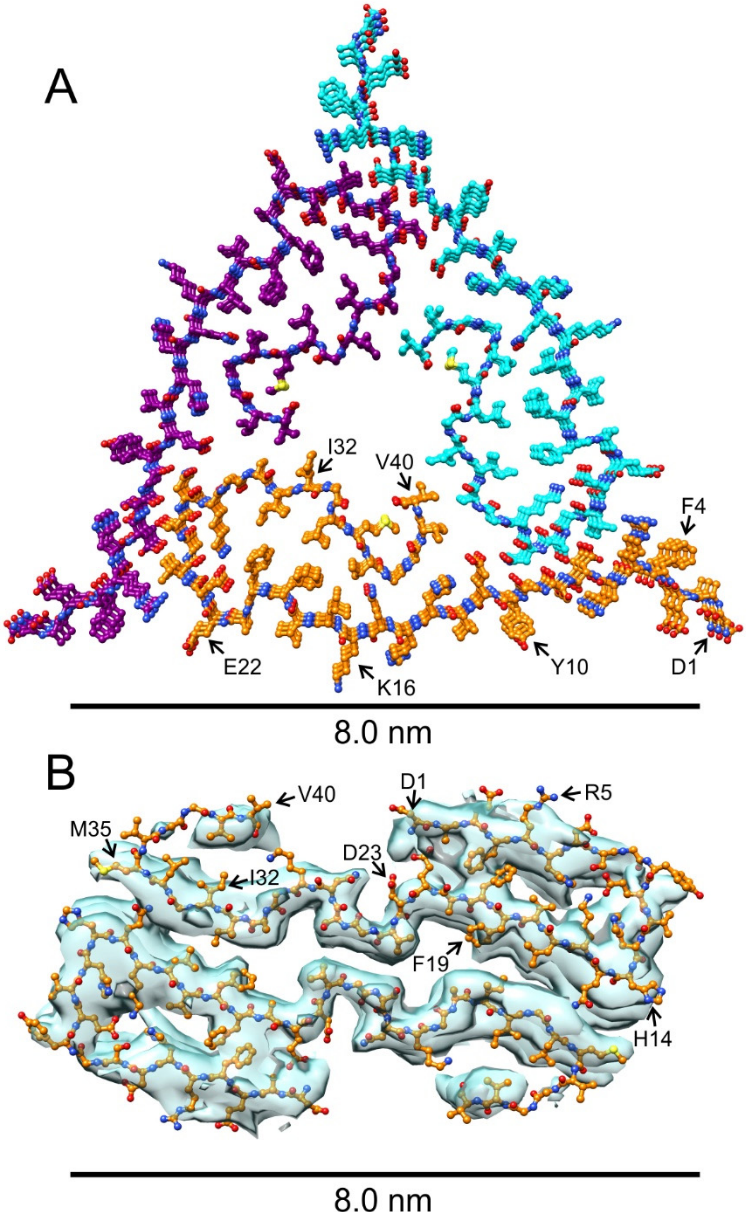


Figure S10: (A) Structural model for brain-derived Aβ40 fibrils from "patient 1" developed from ssNMR data by Lu *et al*. (Protein Data Bank code 2M4J). The fibril is viewed in cross-section, with carbon atoms of the three symmetric cross-β subunits colored orange, cyan, and purple. This model applies to fibrils with a different morphology than those shown in Figs. 1 and S1. (B) Cross-sectional view of a cryoEM density map and molecular model for Aβ40 fibrils isolated from meningeal tissue of AD patients, developed by Kollmer *et al*. (Protein Data Bank code 6SHS, Electron Microscopy Data Bank code 10204). The reported resolution of the density map is 4.4 Å.

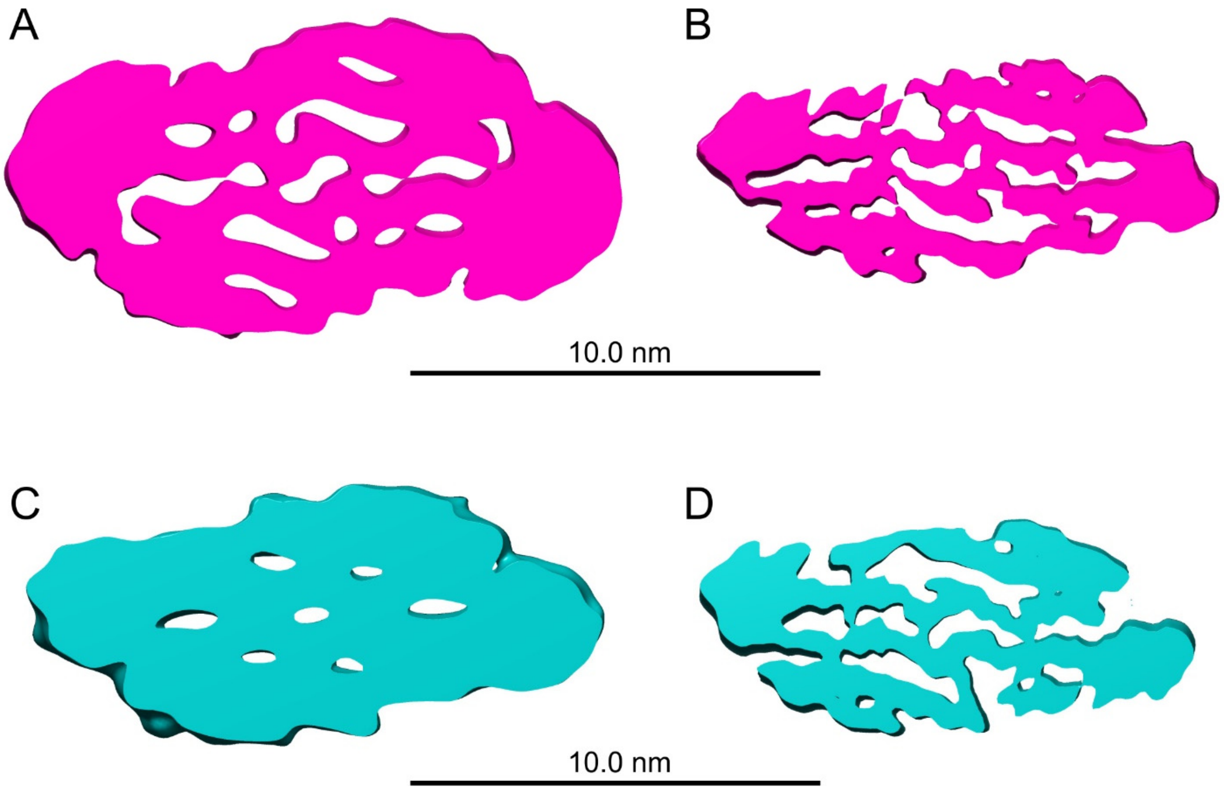


Figure S11: (A) Initial density model for calculations of the cryoEM density map in Fig. 2. This model was obtained from studies of *in vitro* Aβ40 fibrils with morphologies similar to those in Figs. 1 and S1. (B) Density map resulting from the initial stage of 3D classification with unmodified RELION 3.0, described in Step 3 of the Supporting Text. (C) Alternative initial density model obtained from well-resolved 2D class images of brain-derived Aβ40 fibrils. (D) Density map resulting from the initial stage of 3D classification, using the alternative initial model.

Table S1: Parameters for cryoEM data collection, 3D density map reconstruction, and molecular model development.

| **Data Collection** | |
|---|---|
| Magnification | 130,000X |
| Voltage | 300 kV |
| Defocus range | -0.5 µm to -3.0 µm |
| Microscope | Krios |
| Camera | K2 Summit |
| Exposure time | 10 s |
| # movie frames | 50 |
| Total electron dose | 73.5 e/Å$^2$ |
| Pixel size | 1.08 Å (binned from 0.54 Å) |
| Number of images used | 1337 |
| **3D Density Map Reconstruction (EMDB-21501)** | |
| Box size (pixel) | 400 pixels, 432 Å |
| Inter-box distance | 29 Å |
| Number of fibril segments selected | 19790 |
| Number of particles extracted from images | 383,717 |
| Number of particles after 2D classification | 239,937 |
| Number of particles in final density map | 103,462 |
| Resolution | 2.77 Å |
| B-factor | -46.9 Å$^2$ |
| Helical rise | 2.45 Å |
| Helical twist | 179.66° |
| Symmetry | effectively $2_1$ screw symmetry |
| **Atomic Model (PDB 6W0O)** | |
| Number of unique non-hydrogen atoms (residues 10-40) | 234 per molecule |
| RMSD of bonds | 0.01 Å |
| RMSD. of angles | 1.55° |
| Molprobity clash score | 0.72 |
| Favored rotamers | 95% |
| Ramachandran outliers | 0% |
| Ramachandran favored | 95% |
| **Xplor-NIH statistics** | |
| Number of structures in final round of calculations | 60 |
| Total number of violations of ANGL, BOND, CDIH, IMPR, repel, repel14, CDIH, and terms in final structure bundle | 0 |
| Potential energy range of 30 lowest-energy structures (arbitrary units) | 10025.86-10299.90 |
| All-atom RMSD for residues 14-40 of central molecules in final structure bundle | 1.35 Å |
| Backbone RMSD for residues 14-40 of central molecules in final structure bundle | 0.37 Å |

Table S2: Chemical shifts for Aβ40 molecules in the inner cross-β layers of brain-derived Aβ40 fibrils (BMRB code 30731). Shift values are in parts per million (ppm) relatively to sodium trimethylsilylpropane-sulfonate (DSS) for $^{13}C$ and liquid ammonia for $^{15}N$. Uncertainties are approximately ±0.2 ppm for CO and N sites and ±0.1 ppm for other carbon sites.

| | $C_\alpha$ | $C_\beta$ | $C_\gamma$ | $C_\delta$ | $C_\varepsilon$ | CO | N |
|---|---|---|---|---|---|---|---|
| K16 | 54.3 | 34.7 | - | - | - | 174.0 | 123.5 |
| L17 | 54.8 | 44.6 | 30.0 | - | - | 175.0 | 126.5 |
| V18 | 60.2 | 35.1 | 20.8 | - | - | 173.0 | 121.0 |
| F19 | 55.7 | 40.9 | - | - | - | 173.2 | 125.7 |
| F20 | 56.0 | 43.3 | - | - | - | 172.4 | 127.0 |
| A21 | 49.7 | 23.0 | - | - | - | 175.0 | 128.9 |
| E22 | 53.0 | 33.6 | 35.8 | 181.7 | - | 175.5 | 119.9 |
| D23 | 54.0 | 37.4 | 181.5 | - | - | 174.4 | 123.9 |
| V24 | 59.8 | 34.8 | 20.2, 20.8 | - | - | 175.3 | 121.7 |
| G25 | 47.3 | - | - | - | - | 174.3 | 115.6 |
| N27 | 55.4 | - | - | - | - | 174.2 | - |
| K28 | 55.4 | 35.3 | 26.3 | 29.4 | 41.9 | 174.4 | 125.1 |
| G29 | 43.2 | - | - | - | - | 172.0 | 108.3 |
| A30 | 49.8 | 21.5 | - | - | - | 175.1 | 127.4 |
| I31 | 60.8 | 40.0 | 19.0, 28.0 | 13.5 | - | 174.2 | 124.9 |
| I32 | 57.3 | 42.3 | 17.0, 26.6 | 13.7 | - | 176.2 | 124.9 |
| G33 | 48.7 | - | - | - | - | 172.2 | 115.2 |
| L34 | 53.3 | 45.8 | 27.9 | 25.1 | - | 174.1 | 114.7 |
| M35 | 54.0 | 36.8 | 31.7 | - | - | 173.9 | 121.6 |
| V36 | 59.4 | 36.0 | 20.6 | - | - | 173.8 | 123.4 |
| G37 | 44.4 | - | - | - | - | 171.8 | 111.7 |
